# A Normative Framework Dissociates Need and Motivation in Hypothalamic Neurons

**DOI:** 10.1101/2023.10.01.560411

**Authors:** Kyu Sik Kim, Young Hee Lee, Yu-Been Kim, Jong Won Yun, Ha Young Song, Joon Seok Park, Sang-Ho Jung, Jong-Woo Sohn, Ki Woo Kim, HyungGoo R. Kim, Hyung Jin Choi

## Abstract

Physiological needs evoke motivational drives to produce natural behaviours for survival. However, the temporally intertwined dynamics of need and motivation have made it challenging to differentiate these two components in previous experimental paradigms. Based on classic homeostatic theories, we established a normative framework to derive computational models of neural activity and behaviours for need-encoding and motivation-encoding neurons during events that induce predicted gain or loss. We further developed simple and intuitive experimental paradigms that enabled us to distinguish the distinct roles of subpopulations of neurons in the hypothalamus. Our results show that AgRP neurons and LH^LepR^ neurons are consistent with need and motivation, respectively. Our study provides a parsimonious understanding of how distinct hypothalamic neurons separately encode need and motivation to produce adaptive behaviours for maintaining homeostasis.

## Introduction

Physiological needs produce motivational drives to generate natural behaviours crucial for survival (*1-4*). Classic psychological theories have elucidated various aspects of homeostatic need and motivation (*5-8*). In the drive-reduction theory, the deviation of an animal’s internal state from the homeostatic set point is perceived as a “current deficit” (*8*). For survival, animals not only monitor their “current deficit” but also utilise external information to predict future change in state, thereby forecasting a “predicted deficit” (*9-14*). Since this external information helps to predict future change, it acts as “predicted gain (or loss)” which update the “predicted deficit” in a feedforward manner (*14-17*).

Evolutionarily, the predicted-deficit-based strategy is more efficient for survival than the current-deficit-based strategy (*17*). The former enables preemptive preparation for efficient behavioural regulation and avoids overcompensation which might lead to a maladaptive state (*18*). Therefore, the “predicted deficit” serves as the animal’s overall “need”. “Motivation” is produced by need to evoke behaviour and only starts to increase when the animal has accessibility to a specific goal (*19-24*). Therefore, need accumulates into goal-directed motivation when accessibility is provided (*25-28*). When accumulated motivation exceeds a certain threshold, specific behaviours aimed at reducing the “need” are initiated (*29, 30*).

In the context of eating, when an animal perceives its environment as accessible to its goal, need accumulates into food-directed motivation, producing a sequence of natural eating behaviours (*8, 25*). However, no study has yet demonstrated how the need and motivation for food are separately encoded, or how distinct hypothalamic neurons respond sequentially to food-related external events. Agouti-related peptide (AgRP) neurons in the arcuate hypothalamus are the major regulators of eating behaviours (*31*), although their specific psychological role in the homeostatic process is yet to be clarified. Previous studies have demonstrated that AgRP neurons activate to encode food need during fasting, responding to food-anticipatory signals (*32-35*). However, other studies argue that despite the deactivation of AgRP neurons, the residual firing of AgRP neurons directly encodes the food-related motivation responsible initiating behaviours such as searching, foraging, consuming or instrumental responses (*36, 37*). We and other researchers have reported that leptin-receptor neurons in the lateral hypothalamus (LH^LepR^) are activated during food consumption (*38, 39*). Similar to activating AgRP neurons, activation of LH^LepR^ neurons increases eating behaviours (*38*). Collectively, the activation of both AgRP neurons and LH^LepR^ hypothalamic neuron populations evokes eating behaviours which complicate the understanding of their roles as either need or motivation for eating.

Past studies have reported that the concept of homeostasis can be incorporated into a model derived from a normative framework. Recent studies have demonstrated that a combination of experiments and computational models can distinguish (*40, 41*) the temporal dynamics of different psychological variables in the cortex and the limbic systems (*42-44*). Building on these findings, we hypothesised that a computational model, derived from a normative framework, could be applied to the temporal dynamic data of neural activity and behaviours to distinguish the functional roles of different populations of hypothalamic neurons. Therefore, we derived models based on the concept of need and motivation (For detailed framework, see Methods) and applied them to the neural activity of AgRP neurons and LH^LepR^ neurons (LH^LepR^ data from (*38*)), as well as behavioural activity traces during activation of these neurons, in order to distinguish their psychological roles as either need or motivation.

## Result

### Normative Models of Need and Motivation for Food

The food deficit (*D*_*f*_) is a function of homeostatic state (*H*_*t*_) at time t, which can be defined from the deviation of the homeostatic set point as below,

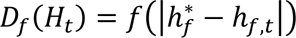

where 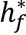 is the homeostatic set point and ℎ_*f*,*t*_ is the homeostatic state at time *t*. For optimal survival, animals use external information to forecast future states. Therefore, a predicted change (*PC*_*f*_(*H*_*t*_)) is defined as a change in future state predicted from external information. A predicted deficit (*PD*_*f*_(*H*_*t*_)) is defined as the sum of current food deficit *D*_*f*_(*H*_*r*_) and all future predicted changes *PC*_*f*_(*H*_*t*_) (*15*) (Fig. 1a, See Methods for detailed description).

**Fig. 1.**
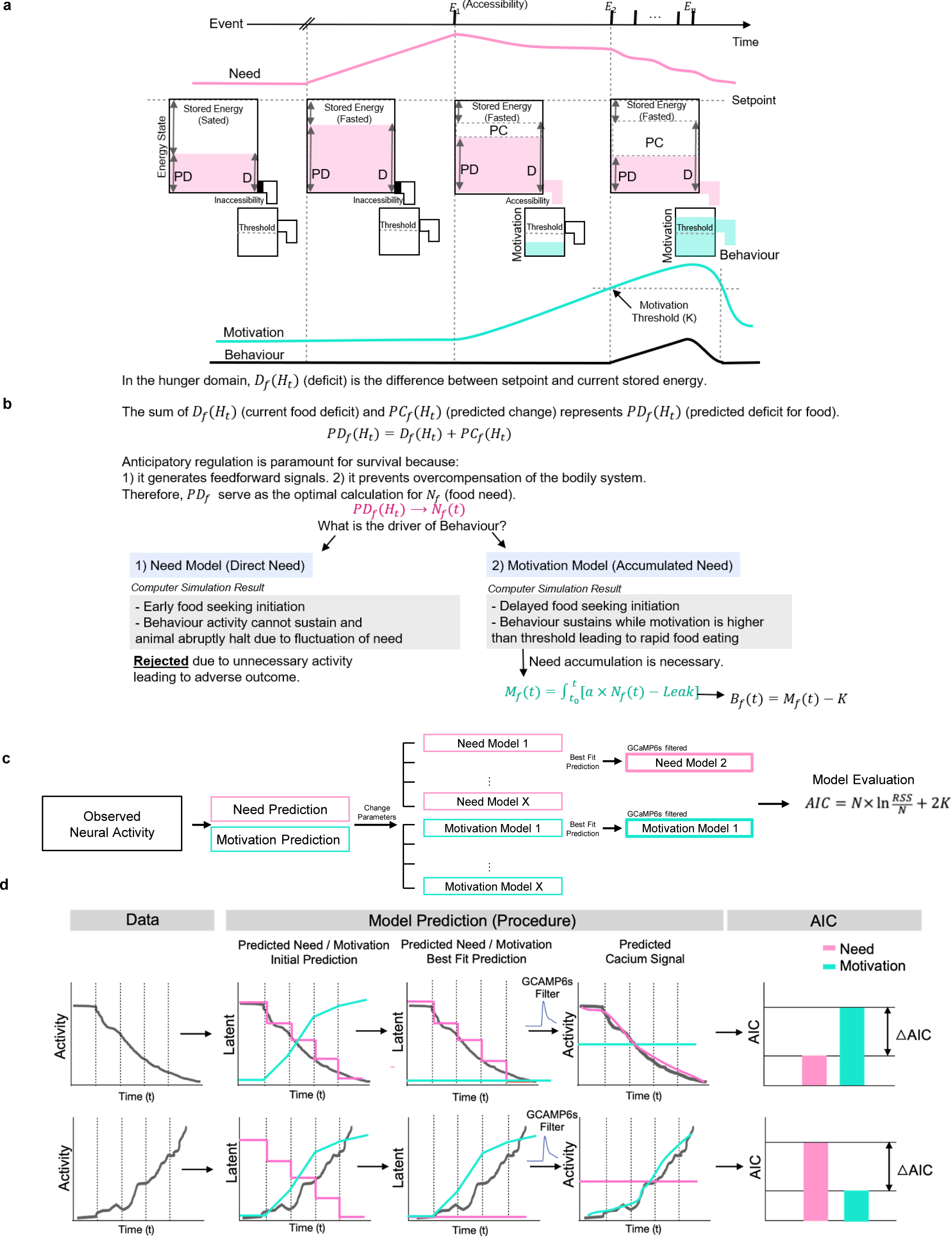
Outline of Normative Framework and Computational Analysis to Dissociate Need and Motivation. **a**, Schematic of the temporal dynamics of need and motivation. **b,** Detailed description of the normative framework for **a**. Simulation result shows that the motivation model(accumulated need) is the driver of behaviour. **c,** Pipeline of the model procedure in need and motivation models. **d,** Schematic of the model-fit procedure. The latent values were obtained and convolved with the GCaMP6s kernel. To determine which model best explains the neuronal activity, Akaike information criterion (AIC) were obtained.

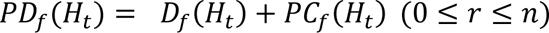

The predicted deficit should serve as the animal’s optimal choice to calculate the current food need *N*_*f*_(*t*) (afterwards stated as need, Fig. 1a,b)

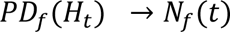

Can *N*_*f*_(*t*) be the main driver of behaviour for an animal? Based on our simulation results the *N*_*f*_(*t*) cannot be a driver of behaviour since it delays food acquisition (Extended Data Fig. 1b,c). Instead, motivation (*M*_*f*_(*t*)), which accumulates from need, serves as a more advantageous policy for generating rapid and efficient behaviour (Supplementary Video 1). We defined the intensity of food motivation as the accumulation of need with the product of integrated affecting factors (*a*) attenuated with a *Leak*:

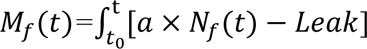

Behaviour *B*_*f*_(*t*) is initiated when the motivation surpasses a threshold (*K*).

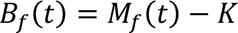

Next, using a model-fitting analysis from our previous study (*42*), we established an analysis pipeline to determine which hypothalamic neurons could be attributed to the need or motivation model, respectively. The need and motivation models were initially defined based on our framework above. For each model, the time course of the latent variable (need or motivation) was defined with experimental events. Then, we obtained a set of best-fit parameters that minimised the difference between predicted and empirical neural responses by employing a model-fit procedure (Fig. 1c,d see Methods).

### Models and Experiments Using Events That Induce Predicted Gain/Loss Dissociate Need and Motivation in Hypothalamic Neurons

Based on our models, we investigated whether the neural activity of hypothalamic neurons best represent the properties of need or motivation. To measure neural activity, AgRP-cre and LepR-cre mice were injected with cre-dependent adeno-associated virus (AAV) carrying GCaMP6s, and an optic fibre was implanted in the ARC or LH, respectively (Fig. 2a-d).

**Fig. 2.**
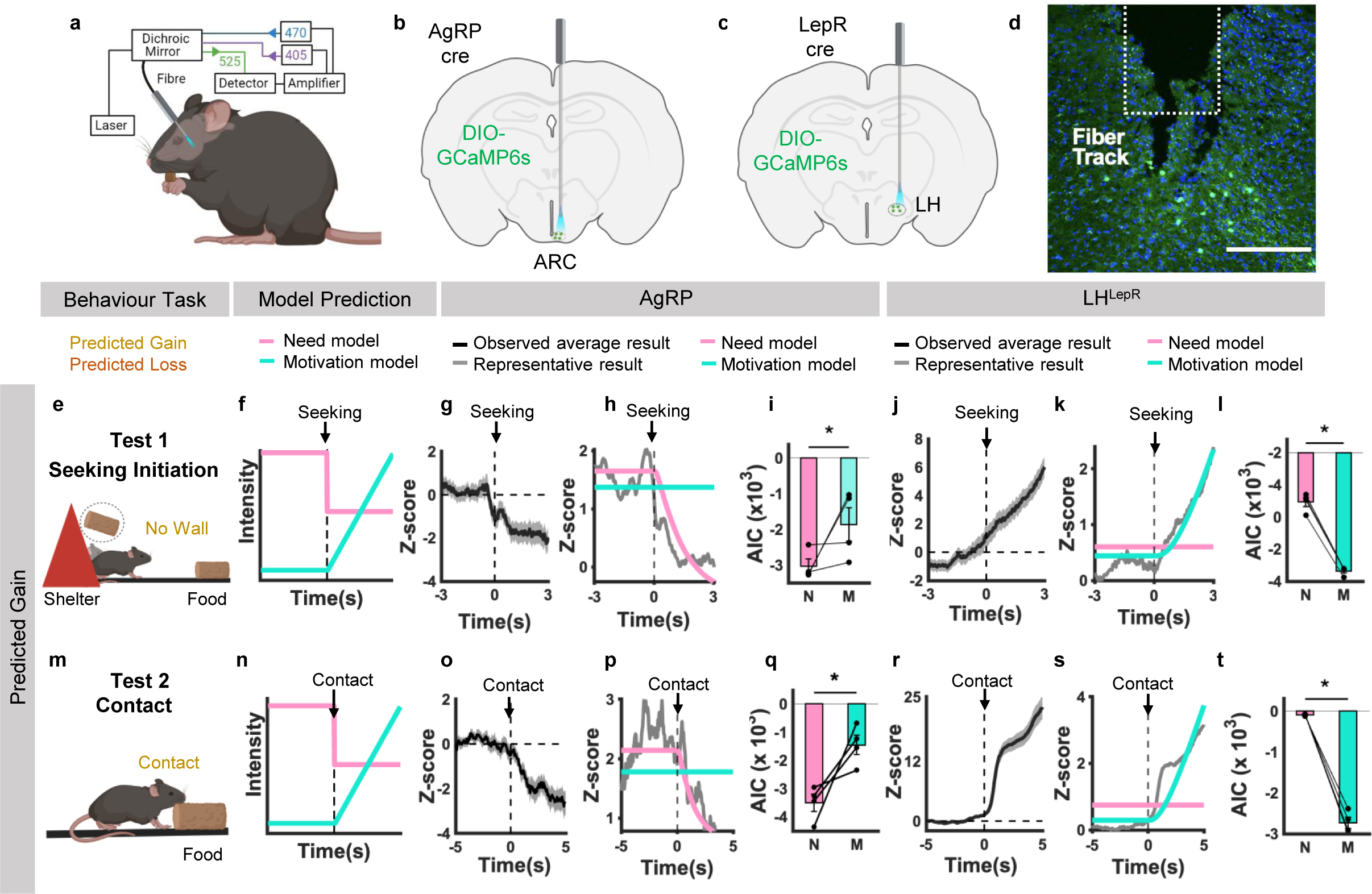
Events That Induce Predicted Gain Dissociate AgRP and LH^LepR^ Neurons as Need and Motivation, Respectively. **a**, Schematic of photometry experiment design **b, c,** Schematic of viral injection into ARC **b** and LH **c. d,** Representative image validates GCaMP6s expression in LH^LepR^ neurons. **e, m,** Schematic of behavioural paradigm and event. **e**, Predicted Gain Test 1, **m**, Predicted Gain Test 2. **f, n,** Schematic of model prediction of need (pink) and motivation (turquoise) intensity in each test. Dotted line indicate the behaviour of interest in each test. **f,** Seeking initiation moment, **n,** Contact moment. **g, o, j, r,** Average trace of Z-score from all trials from individual mice for each test. Vertical l dotted line at 0s indicate the moment of behaviour of interest. Horizontal dotted lines indicate when Z-score is 0. **g, o,** Neural activity from AgRP neurons in each test. **j, r,** Neural activity from LH^LepR^ in each test. **h, p, k, s,** Model fitting result between neural activity data (Grey) and need neural activity model (pink) or motivation neural activity model (turquoise) for each test. **h, p,** Model fitting of AgRP neural activity in each test. **k, s,** Model fitting of LH^LepR^ neural activity in each test. Vertical dotted line at 0s indicate the moment of behaviour of interest. i**, q,** Quantification of AIC from AgRP neural activity and need neural activity model (pink) or motivation neural activity model (turquoise) in each test. **l, t,** Quantification of AIC from LH^LepR^ neural activity and need neural activity model (pink) or motivation neural activity model (turquoise) in each test. **e, f, g, h, I, j, k, l**, Data from Predicted Gain Test 1 (AgRP, LH^LepR^ N= 4,4 Trials= 46,67). **m, n, o, p, q, r, s, t,** Data from Predicted Gain Test 2 (AgRP, LH^LepR^ N= 4,4 Trials= 32,32). Data are mean ± s.e.m. See Supplementary Table. 1 for statistics. Source data are provided as a source data file. The schematics in **a** and **m** were created using BioRender.

We first conducted two experiments in fasted mice to identify which hypothalamic neurons encode need or motivation. These experiments were specifically designed to evoke events that induce predicted gain: 1) Seeking initiation (predicts food discovery), 2) Contact (predicts nutrient absorption). In the seeking initiation experiment, according to our models, there is a decrease in need and an increase in motivation at the moment of voluntary seeking initiation event (Fig. 2e-f). Notably, in our neural recordings, the AgRP neural activity started to decrease at the voluntary seeking initiation event (Fig. 2g), while the neural activity of LH^LepR^ started to increase (Fig. 2j). Additionally, we observed that the neural onset of both AgRP neurons (mean = −6.656s) and LH^LepR^ neurons (mean = −5.623s) preceded the voluntary seeking initiation, which suggests neural activity change is the cause of behaviour (Extended Data Fig. 3). Comparisons of the goodness-of-fit revealed that the AgRP and LH^LepR^ neural activity were significantly more consistent with the need and motivation models, respectively (Fig. 2h,i,k,l). In the contact experiment, according to our models, there is a decrease in need and an increase in motivation at the moment of food contact event (Fig. 2m,n). In our neural recordings, AgRP neural activity started to decrease at the food contact (Fig. 2o), while the neural activity of LH^LepR^ started to increase (Fig. 2r), which is consistent with the need model and motivation model, respectively (Fig. 2p,q,s,t).

To further distinguish the identity of hypothalamic neurons, we investigated neural activity during events that induce predicted loss. 1) Inaccessibility (predicts prolonged starvation), 2) Abandon (predicts food loss). In the inaccessibility experiment, according to our models, the predicted loss produced an increase in need. However, the motivation will not change because the inaccessibility blocks access to the goal (Fig. 3a,b). Interestingly, the neural activity of AgRP increased at the inaccessibility event, whereas the neural activity of LH^LepR^ remained stable (Fig. 3c,f). These results were consistent with the need model and motivation model, respectively (Fig. 3d,e,g,h). In the abandon experiment, according to our models, predicted loss produce an increase in need and a decrease in motivation (Fig. 3i,j). Consistent with the need model, the neural activity of AgRP increased at the moment of eating abandon and showed sustained activity (Fig. 3k-m). The neural activity of LH^LepR^ was inactivated, consistent with the motivation model (Fig. 3n-p). Similarly, model-fitting results of individual trials from each mouse confirmed the same conclusion in a similar behaviour paradigm upon voluntarily abandoning eating after seeking behaviour (Extended Data Fig. 4).

**Fig. 3.**
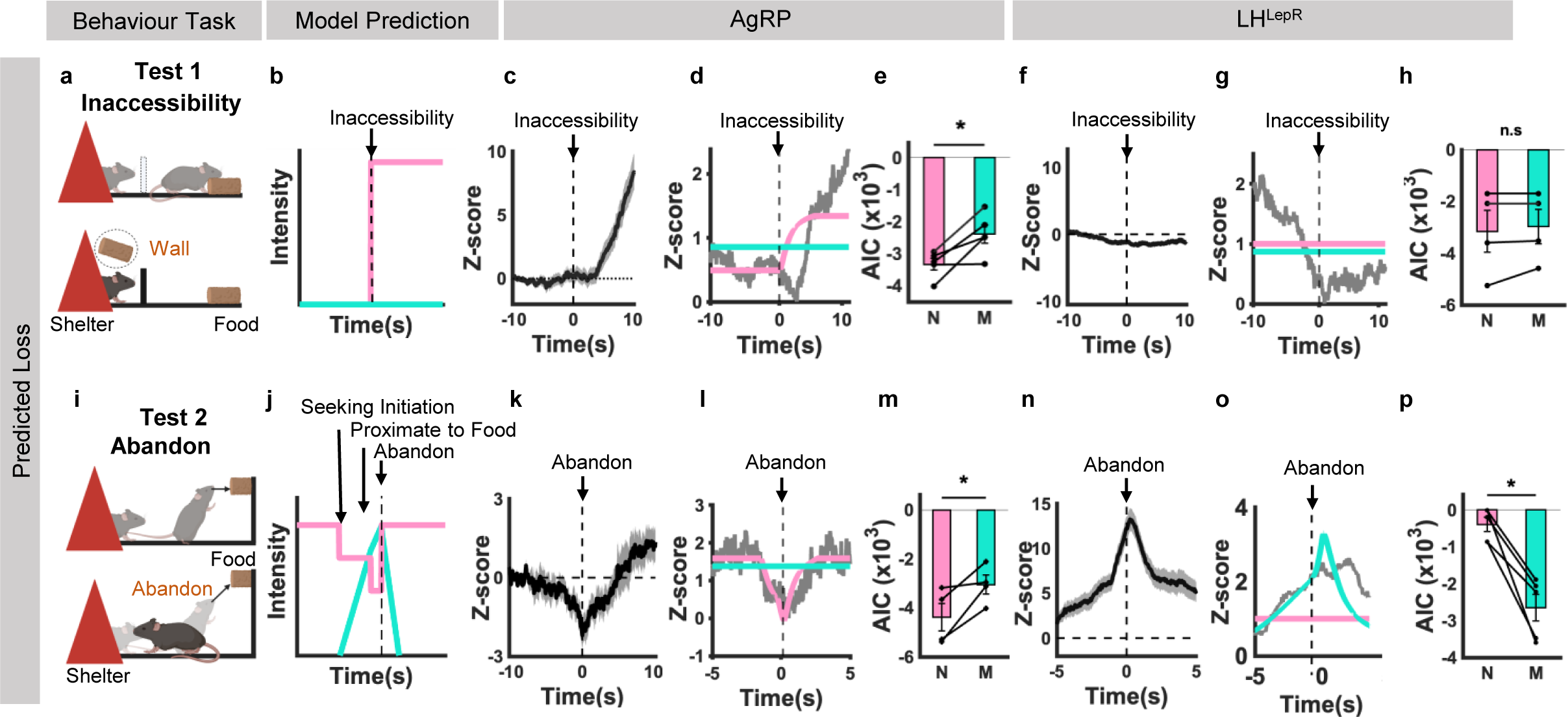
Events That Induce Predicted Loss Dissociate AgRP and LH^LepR^ Neurons as Need and Motivation, Respectively. **a, i**, Schematic of behavioural paradigm and event. **a,** Predicted Loss Test 1 **i,** Predicted Loss Test 2. **b, j,** Schematic of model prediction of need (pink) and motivation (turquoise) intensity in each test. Dotted line indicate the behaviour of interest in each test. **b,** Inaccessibility moment, **j,** Abandon moment. **c, k, f, n,** Average trace of Z-score from all trials from individual mice for each test. Vertical l dotted line at 0s indicate the moment of behaviour of interest. Horizontal dotted lines indicate when Z-score is 0. **c, k,** Neural activity from AgRP neurons in each test. **f, n,** Neural activity from LH^LepR^ in each test. **d, l, g, o,** Model fitting result between neural activity data (Grey) and need neural activity model (pink) or motivation neural activity model (turquoise) for each test. **d, l,** Model fitting of AgRP neural activity in each test. **g, o,** Model fitting of LH^LepR^ neural activity in each test. Vertical dotted line at 0s indicate the moment of behaviour of interest. **e, m,** Quantification of AIC from AgRP neural activity and need neural activity model (pink) or motivation neural activity model (turquoise) in each test. **h, p,** Quantification of AIC from LH^LepR^ neural activity and need neural activity model (pink) or motivation neural activity model (turquoise) in each test. **a, b, c, d, e, f, g, h** Data from Predicted Loss Test 1 (AgRP, LH^LepR^ N= 5,4 Trials= 75,60). **i, j, k, l, m, n, o, p,** Data from Predicted Loss Test 2 (AgRP, LH^LepR^ N= 4,5 Trials= 20,17). Data are mean ± s.e.m. See Supplementary Table. 1 for statistics. Source data are provided as a source data file. The schematics in **a** and **i** were created using BioRender.

The dissociation of distinct neural dynamics was further examined using unbiased dimensionality reduction methods. Principal component analysis (PCA), incorporating both types of hypothalamic neurons, further showed that AgRP and LH^LepR^ neurons exhibit different temporal dynamic trajectories during multi-predicted gain behaviours (Fig. 4a,b). These neural trajectories were also distinctly separated in a t-SNE dimension (Fig. 4c). Furthermore, a nonlinear analysis method that unpacks neural dynamics in a high-performance latent space (CEBRA) (*45*) also demonstrated different embeddings (Extended Data Fig. 2, Supplementary Movie 2). These results further demonstrate that even without assuming any theory-driven models, the population responses form distinguishable dynamics on their own.

**Fig. 4.**
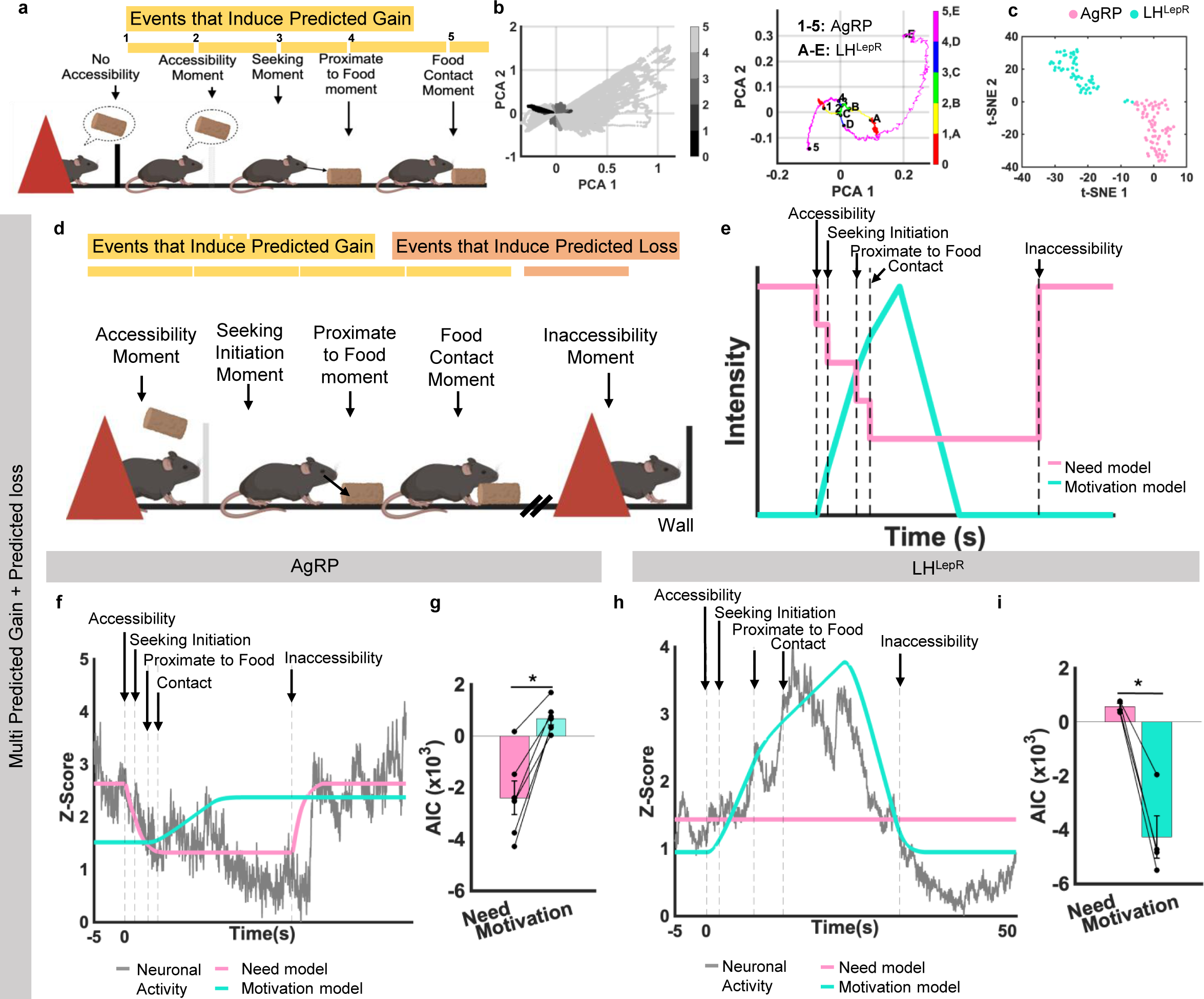
Multi-Events That Induce Predicted Gain/Loss and Theory-Driven Models Dissociate AgRP and LH^LepR^ Neurons as Need and Motivation, Respectively. **a,** Schematic of behavioural paradigm of multi-predicted gain. **b,** PCA analysis of all neural trajectories (left) and average neural trajectories of AgRP neurons (1-5) and LH^LepR^ neurons (A-E), 0, Start of trial, 1, A, Accessibility, 2, B, Seeking initiation, 3, C, Proximate to food, 4, D, Contact, 5, E, End of trial (right). **c,** t-SNE analysis of all neural trajectories. **d,** Schematic of behavioural paradigm of multi-predicted gain/loss test **e,** Schematic of model prediction of need (pink) and motivation (turquoise) intensity. **f, h,** Representative model fitting result between neural activity data (normalized Z-score, grey) and need neural activity model (pink) or motivation neural activity model (turquoise) from an individual trial. **f,** Neural activity from AgRP neurons (N= 6, Trials= 56), **h,** Neural activity from LH^LepR^ neurons (N = 4, Trials = 50). **g, i,** Quantification of AIC between neural activity and need neural activity model (pink) or motivation neural activity model (turquoise). **g,** AIC quantification from AgRP neurons (N = 6), **i,** AIC quantification from LH^LepR^ neurons (N = 4). Data are mean ± s.e.m. See Supplementary Table. 1 for statistics. Source data are provided as a source data file. The schematics in **a** and **d** was created using BioRender.

Natural eating behaviours involve multiple sequential events that induce consecutive predicted changes in energy homeostasis. To validate our findings in more naturalistic eating behaviours, which contain both predicted gain and loss, we conducted a multi-predicted gain/loss test (Fig. 4d). After events that induce multi-predicted gain, fasted mice were subjected to an event that induce predicted loss (inaccessibility). We predicted that need would decrease whenever a predicted gain occurred and would show a rebound when inaccessibility occurred. Conversely, motivation would start to increase at the accessibility moment and then gradually decline after consumption. According to our models, at the inaccessibility event, the need increases, whereas motivation remains unchanged (Fig. 4e). In our neural recordings, AgRP and LH^LepR^ neural activities were consistent with the need and motivation model, respectively (Fig. 4f-i and Supplementary Movie 3). We further examined the neural activities using individual trials. Due to the self-paced nature of the task, the timing of these sequential events varied across trials, and responses were noisier. However, our model-fit results using individual trials clearly preferred one model over the other (Extended Data Fig. 5).

We further examined whether neural activity reflects transitions of states through multiple events instead of a single event, since AgRP neurons have primarily been tested with single events that induce predicted gain (cue or sensory detection) (*36*). As expected, the best fit for AgRP neural activity was the need model, which decreased at each predicted gain rather than single-event decrease models (Extended Data Fig. 6). Similarly, for LH^LepR^ neurons, the best fit was the multi-event motivation model (Extended Data Fig. 7). These results indicate that when animals learn a sequence of food-predicting events, the neurons predicting food intake show a cascade of shifts, rather than one big shift, as the animals perform the task. Collectively, these results demonstrate that the neural activity of AgRP and LH^LepR^ is temporally dissociable and consistent with need and motivation, respectively.

### Theory-Driven Models Explain Dynamics of Behaviours Evoked by Activation of Hypothalamic Neurons

Next, we investigated whether the manipulation of these neurons affects behaviours, similar to the predictions of our models. To do so, we injected channelrhodopsin (ChR2) virus and implanted optic fibres in the ARC or LH (Fig. 5a,b) and tested ad-libitum mice in an eating-evoking test, activating target neurons for 10 s (Fig. 5c).

**Fig. 5.**
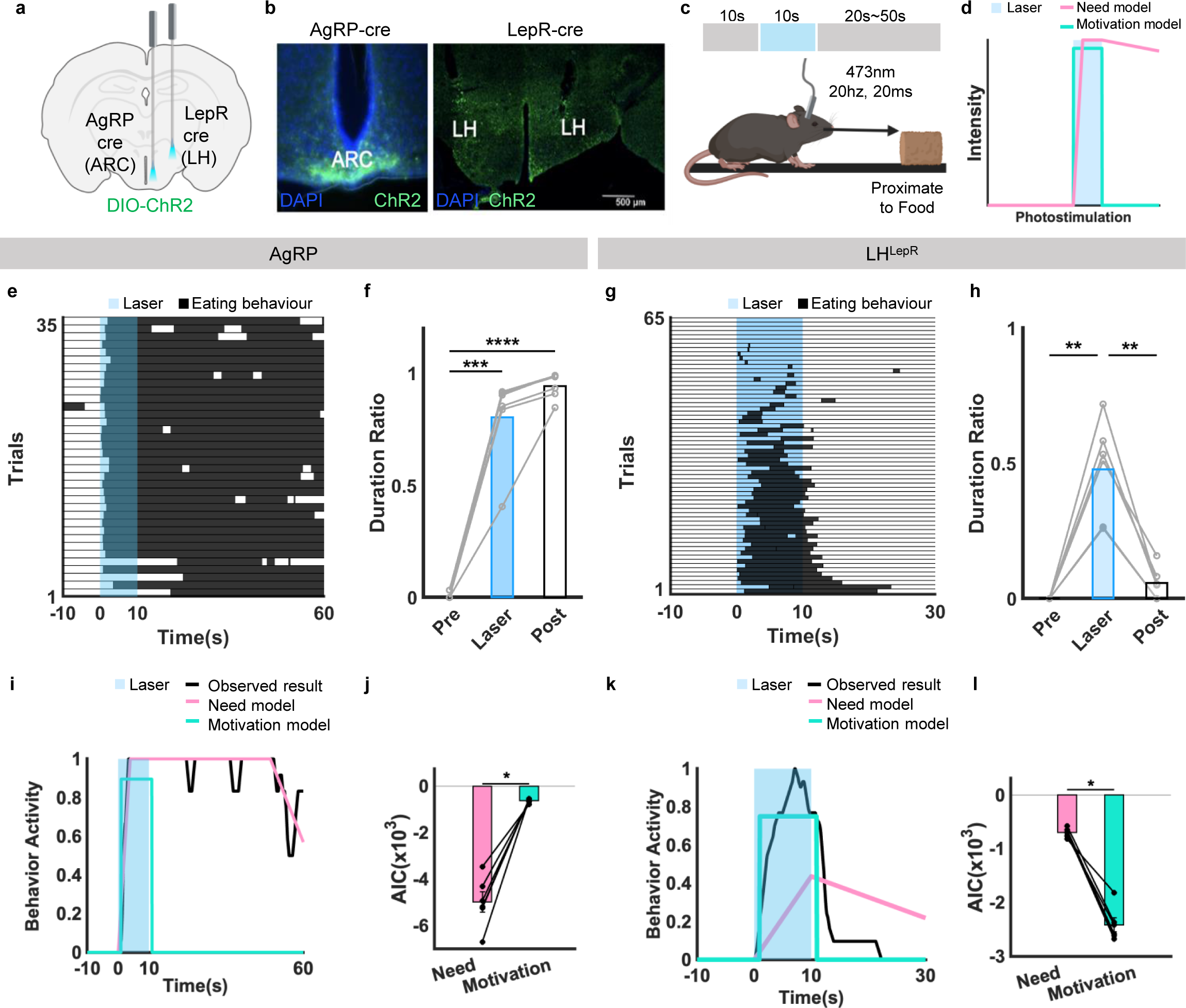
Theory-Driven Models Dissociate Behaviour Activity Evoked from AgRP and LH^LepR^ Neural Activation as Need Evoked Behaviour and Motivation Evoked Behaviour, Respectively. **a**, Schematic of virus injection into the target site. **b,** Representative image validates ChR2 expression in AgRP and LH^LepR^ neurons. **c,** Schematic of behavioural paradigm and event. **d,** Schematic of model prediction of behaviour activity evoked from need (pink) and motivation (turquoise) neuronal activation. Duration of optogenetic stimulation coloured in blue. **e,** Raster plot of eating behaviour (black) shown for each individual trial from all AgRP mice. Duration of optogenetic stimulation shown in blue (N = 6, Trials = 36). **f**, Behaviour quantification of behaviour duration ratio from **e**. **g,** Raster plot of eating behaviour (black) shown for each individual trial from LH^LepR^ mice (N = 4, Trials = 65). Duration of optogenetic stimulation shown in blue. **h,** Behaviour quantification of behaviour duration ratio from **g**. **i, k,** Model fitting of behavior activity between behavior activity data from AgRP or LH^LepR^ neuronal activation (black) and behavior activity prediction from need neuron activation (pink) or motivation neuron activation (turquoise). **i,** Behaviour activity from AgRP Neuronal activation (1 mice, 6 trials). **k,** Behaviour activity from LH^LepR^ Neuronal activation (1 mice, 10 trials). **j, l,** Quantification of AIC between behaviour activity from AgRP or LH^LepR^ neuronal activation and need behaviour activity model (pink) or motivation behaviour activity model (turquoise). **j,** Quantification of AIC from AgRP neuronal behaviour activation (N=6). **l,** Quantification of AIC from LH^LepR^ neuronal behaviour activation (N=6). Data are mean ± s.e.m. See Supplementary Table. 1 for statistics. Source data are provided as a source data file. The schematics in a and c were created using BioRender.

Our models predict that optogenetic activation of need and motivation neurons would result in different temporal dynamics of behaviour. If need neurons were activated, the need would rapidly surge to its highest intensity and then gradually accumulate to generate motivation. Once the accumulated motivation surpasses the behaviour initiation threshold, it would produce the behaviour. When need neuron activation terminates, the intensity of the need would return to null, and motivation would gradually decrease. The behaviour would continue until motivation decreases below the behaviour threshold. On the other hand, if the motivation neurons were activated, the motivation level would surge to its highest intensity, immediately surpassing the behaviour threshold to produce the behaviour. When motivation neuron activation terminates, the motivation would drop to its lowest intensity, leading to the immediate cessation of the behaviour activity. Therefore, the AgRP and LH^LepR^ neurons can be predicted to exhibit different eating behaviour activity dynamics (Fig. 5d).

As predicted, the activation of AgRP neurons showed significant sustained behaviour activity even after the termination of activation (Fig. 5e,f) while manipulating LH^LepR^ neurons showed an immediate cessation of behaviour activity upon deactivation (Fig. 5g,h). When we fitted the temporal dynamics of behavioural responses evoked by activating hypothalamic neurons with our models, AgRP and LH^LepR^ behaviour activity was indeed significantly closer to the need model and motivation model, respectively (Fig. 5i-l and Supplementary Movie 4). Collectively, our results disambiguate the roles of hypothalamic neurons, dissociating AgRP neurons as need neurons, and LH^LepR^ neurons as motivation neurons.

## Discussion

In this study, we demonstrated that activities of hypothalamic neurons can be explained by the models of need and motivation in eating behaviours (Extended Data Fig. 8 and Supplementary Movie 5). By incorporating significant events, we were able to differentiate between need and motivation in recorded neural activities. Through model-fit analyses, we revealed that the AgRP and LH^LepR^ neurons encode need and motivation, respectively. Moreover, by manipulating the two types of neuron, we observed distinct temporal patterns in eating behaviours and our behaviour activity models accurately predicted these differences. Collectively, our findings shed light on the neural basis of distinct psychological constructs in eating behaviour.

### Normative Framework to Investigate the Neural Substrate of Eating Behaviours

Normative modeling analysis has proven to be a valuable approach for understanding individuals with diverse variations and features within human groups (*46*). Despite the aid of computational methods in a sophisticated experimental design to observe natural eating behaviour, determining the homeostatic role of hypothalamic neurons that evoke eating behaviour has remained challenging (*39*). In our study, we demonstrated that the functions of AgRP and LH^LepR^ neurons can be normatively explained, using models derived from recent theories of homeostasis. Our study employed the normative framework to precisely elucidate the neural substrates of need and motivation within the heterogeneous hypothalamic neuronal populations, namely, AgRP and LH^LepR^ neurons. This normative framework provided a temporal dynamic understanding of how these neurons function at a comprehensible level.

### Computational Approaches for Understanding the Neural Representation of Psychological Components

The original drive-reduction theory lacked quantitative details, making it challenging to identify neural substrates of psychological latent variables in the brain. Moreover, the smooth internal transition of need and motivation in natural conditions further complicated their identification. To address these challenges. various computational methods have been developed to identify low-dimensional dynamics of latent variables from high-dimensional data with minimal assumptions and latent variables can be explicitly defined based on theoretical frameworks. The models of these latent variables can be valuable in dissociating different behavioural models when assessing experimental manipulations (accessibility). In our study, we examined low-dimensional components and embeddings to confirm that representative features differ between the two populations. Furthermore, we investigated the nature of these populations using theory-driven models by directly modeling latent variables and adjusting the slow dynamics of measured signals (Fig 1d, GCaMP6s filter). Our model successfully accounted for both trial-averaged activities and trial-to-trial activities.

One may argue, however, that these analyses show correlations between latent variables and neural activities, thus having some limitation in their explanatory power. To addressing this issue, our optogenetic manipulations demonstrated that the two neural populations indeed influenced behaviours with temporal characteristics predicted by the exact model used in the model fit analyses. The high temporal resolution of optogenetic manipulation allowed us to confirm the critical operation in our model, the accumulation of need. Collectively, our integrative approach can be further applied to other hypothalamic neurons, encoding different homeostatic need and motivation features, (thermoregulation, thirst, and osmolarity) to uncover hidden latent features and explain behaviours.

In an evolutionary context, our computer simulation sheds light on the adaptation where organisms have evolved to engage in the process of eating by accumulating need to produce motivation. This evolutionarily driven shift toward motivation-guided behaviour highlights the intricate interplay between physiological need and the motivation process, ultimately shaping survival strategies.

### Deciphering Temporal Neural Activity in Hypothalamic Neurons

Interestingly, during events that induce predicted loss (inaccessibility; abandon moments), AgRP neural activity showed rapid increase within seconds. This represents the first evidence that AgRP neural activity rapidly increases upon cognitive events in adult mice, consistent with the results observed in neonatal mice during events that induce predicted loss (isolation from the nest) (*47*). Along with their responding to voluntary food-seeking initiation, the predictive nature of AgRP neurons suggests that more cognitive components, may be involved in the hypothalamic and subcortical processes related to hunger.

Our optogenetic results demonstrate that the temporal dynamics of behaviour activity induced by AgRP neural activation are consistent with need-induced behaviour activity models. Previous optogenetic studies on AgRP neurons perplexed temporal dynamics of neural activity of behaviour outcomes due to averaging methods (*31, 37*). Our study, along with previous studies, showed that mice exhibit sustained eating behaviours after terminating the AgRP neural activation (*48*). Furthermore, since reaching the motivation threshold requires a sufficient accumulation of need, our models explain why animals exhibit substantial latency in initiating behaviour after food need arises (*49*). Collectively, our model of need accumulation to generate motivation can explain the origin of sustained behaviours and the latency of AgRP neural activity.

Previous studies have not fully accounted for the multitude of dynamic events that can influence the activity of LH^LepR^ neurons (*39, 50*). Past fragmented studies demonstrated that LH^LepR^ neurons reinforce lever pressing, possess rewarding properties (*50*), and initiation of seeking or consummatory behaviours for food (*38*). In our experimental paradigm, LH^LepR^ neural activity increased at accessibility moment and decreased at inaccessibility moment. Our optogenetics results comprehensively demonstrate that the immediate incline and decline of behaviour activity is time-locked to stimulation, consistent with our models. Collectively, these results demonstrate that LH^LepR^ neurons encode food-directed motivation.

Overall, our work unravels how the brain rapidly orchestrates intricate psychological components of homeostatic behaviours to ensure efficient survival.

## Methods

### Animals

All experimental protocols were approved by the Seoul National University Institutional Animal Care and Use Committee and were performed as per the health guidelines for the care and use of laboratory animals from the Seoul National University. Mice were housed on a 08:00 to 20:00 light cycle, with standard mouse chow and water provided ad-libitum, unless otherwise noted. Behavioural tests were performed in the behaviour chamber during the light cycle. Adult male mice (at least 8 weeks old) were used for all behavioural experiments from the following Cre recombinase-expressing mouse lines: LepR-Cre, JAX stock no.008320, a gift from Ki Woo Kim, Yonsei University; AgRP-IRES-Cre JAX stock no. 012899 a gift from Jong-Woo Sohn, Korea Advanced Institute of Science and Technology, were used.

### Stereotaxic Virus Injection

Cre recombinase-expressing mouse lines and Cre-dependent AAV vectors were used. AgRP-ires-Cre mice were injected with the virus to study arcuate nucleus (ARC), and LepR-Cre mice were injected with the virus to study LH. The mice were anesthetised with xylazine (20 mg/kg) and ketamine (120 mg/kg) in saline and placed in a stereotaxic apparatus (KOPF or Stoelting).

For calcium imaging experiments (fibre photometry), a pulled glass pipette was inserted into the target site of the ARC (300 nl total) at the coordinates AP, −1.3 mm; ML, ±0.2 mm; DV, 5.85 mm, from the bregma for AgRP mice, and the LH (400 nl total), at the coordinates AP, −1.5 mm; ML, ±0.9 mm; DV, 5.25 mm, from the bregma for LepR mice. After 10 min, the GCaMP6 virus (AAV1.Syn.Flex.GCaMP6s.WPRE.SV40, Addgene; titre 1.45×10^13^ genome copies per ml) was injected unilaterally for 10 min using a micromanipulator (Nanoliter 2010). The glass pipette was kept at the target site for 10 min following infusion; it was then withdrawn gradually. For optogenetic experiments, opsin virus (AAV5.EF1a.DIO.hChR2 (H134R). EYFP, Addgene; titre 2.4×10^13^ genome copies per ml) was unilaterally injected into the ARC (300 nl; AP, −1.3 mm; ML, ±0.2 mm; DV, 5.85 mm, from the bregma) for AgRP-ires-mice and bilaterally injected into the LH (400 nl; AP, −1.5 mm; ML, ±0.9 mm; DV, 5.25 mm, from the bregma) for LepR mice.

### Optical Fibre Insertion

An optical fibre was inserted on the same day of injecting the virus. The optical fibre was implanted 30 min after injecting the virus to prevent backflow. For fibre photometry experiments, a ferrule-capped optical cannula (400 µm core, NA 0.57, Doric Lenses, MF2.5, 400/430–0.57) was placed unilaterally 0–50 µm above the site of virus injection as previously described; it was attached to the skull with Metabond cement (C&B Super Bond). For optogenetic activation of AgRP neurons, a unilateral optical fibre (200 µm core, NA 0.37, Doric Lenses, ZF1.25_FLT) was implanted 100 µm above the ARC injection site and secured to the skull with Metabond cement. For optogenetic manipulation of the LH^LepR^ neurons, optic fibre was implanted bilaterally 100–500 µm above the site of LH injection at a 10° angle from the vertical in the lateral to the medial direction; it was affixed to the skull with Metabond cement. Before and after the surgery, dexamethasone, ketoprofen, and cefazolin were administered for postoperative care. All mice were recovered in their cages for at least two weeks before conducting the behavioural experiments. The mice were handled for five days to relieve stress and acclimated to the behaviour chamber for 30 min before testing.

### Behaviour tests Animal conditioning

Before the experiments, mice were habituated to the experimental chambers, and fibre handling was conducted for three days. Mice were fasted 80%–90% of the body weight in the ad-libitum state.

### Predicted Gain Test 1

A shelter was placed and an electrical shock was given as punishment in a square chamber, as previously reported (*38*). All reward-associated cues (e.g., visual or sound cues) were eliminated to measure voluntary seeking behaviour initiation. During the conditioning sessions, mice received a chocolate-flavoured snack at the end of the corridor. During the test session, the shock was excluded. Additionally, the moment when the mouse’s whole body emerged out of the shelter (seeking initiation) was analysed.

### Predicted Gain Test 2

Fasted mice received chocolate-flavoured snacks in the L-shaped chamber (60 cm × 8.5 cm) during each trial. The moment when the mice made physical contact with the food was analysed.

### Predicted Loss Test 1

Fasted mice received a chocolate-flavoured snack at the edge of an L-shaped chamber (60 cm × 8.5 cm) with a shelter (6 cm × 12 cm × 18 cm triangle box). During conditioning sessions (three days), each trial started when a door was removed (“accessibility moment”) with scheduled timing from the experimenter. During the test sessions, the mice were either chased or pushed by force into the shelter by the experimenter after receiving a chocolate-flavoured snack. The door was reinstalled in front of the entrance of the shelter to deprive accessibility to the chocolate-flavoured snacks.

### Predicted Loss Test 2

A chocolate-flavoured snack was placed in the tray on one side of the wall. During conditioning sessions, the fasted mice consumed the food in the tray, placed at an obtainable height of 8 cm. During test sessions, fasted mice initiated eating behaviour, rearing toward the visible food; however, the mice eventually abandoned eating behaviours when they realised that the food tray had an unobtainable height of 11 cm. The moment the mice voluntarily abandoned eating behaviours to the hanging food was analysed.

### Predicted Loss Test 3

A food cue (vertical stripe) and a no-food cue (horizontal stripe) were randomly presented at the edge of an L-shaped chamber. During conditioning, mice received chocolate-flavoured snacks only when the food cue was presented. The success rate was recorded during training until it reached 80%, as reported previously (*38*). During the experiment, although the fasted mice initiated seeking after the food cue was presented, eventually they abandoned eating voluntarily, on realising that food was not available.

### The Multi-Predicted Gain test

Conditioning sessions (days 1–2) were performed for 15 trials in each day to provide sufficient experience to the mice for learning about the location of a chocolate-flavoured snack. The test session (day 3) was also performed for 15 trials. Each trial started when a door was removed (“accessibility moment”) with the scheduled timing from the experimenter. “Seeking initiation” was defined when the mice started emerging out of the shelter. ‘‘Proximate to food” was defined when mice arrived at the top of the bridge. “Food contact” was defined as the moment when the mice came in physical contact with the food.

### The Multi-Predicted Gain/Loss test

Fasted mice received a chocolate-flavoured snack in the same chamber of the multi-predicted gain test. The experimenter closed the door after the end of the multi-gain events to measure neural activity predicted loss when loss occured.

### Eating-Evoking Test

Ad-libitum mice were caged in a small cylindrical structure (10 cm diameter × 15 cm height) and habituated for 5 min before laser stimulation. When the mice were proximate to food (i.e., when the mice were indisputably headed toward food), they were randomly given laser stimulation for 10 s. The LH^LepR^ neurons were stimulated for 10 s after a 1 min interval between each trial, and the AgRP neurons were given a wash-out period of at least 1 h after each trial to minimise the sustained effect of the feeding behaviour (*48*).

### Neural Activity Model

#### Preprocessing of Data and Kernel for Model-Fitting

In this study, the preprocessing included fitting the fluorometric data to the first event point, performing Z-scoring, conditional averaging, and up-sampling the data to 100 Hz. Single-trial analyses were performed similarly, but without conditional averaging.

To obtain the kernel of GCaMP6s and compare the predicted latent to the photometric data of the neurons, a single-pulse response filter was used (*51*). In addition, an offset was introduced to address the effect of negative values in the Z-scored signal. A sharp increase occurred at the beginning of the predicted data due to the convolution operation using the GCamp6s kernel. For the multi-predicted gain test, we used the last 5s out of the 10s initial inaccessibility duration. For all other tasks, we added the first 125 samples before trial start with the average value of the first 50 samples after start.

### Model-Fitting Procedure

Need and motivation models predict activity of neurons based on the specific events of each experiment. The fitting equations used for each experiment are explained in the following sections, from which the latent values were obtained and convolved with the GCaMP6s kernel to predict the actual data. The predicted values were then compared to the actual data to obtain the optimal set of parameters with the lowest RMSE. In the single-trial analysis, we used neural response and predictions from events that vary across trials. The same set of parameters was used for performing minimisation of RMSE for each experimental session.

An initial set of 100 parameters was randomly provided for executing the aforementioned process. MATLAB’s fmincon function was used to find the optimal parameters with the lowest RMSE. Finally, the Akaike information criterion (AIC) (*52*) was obtained to compare the performance of models and determine which model best explained the neural activity.

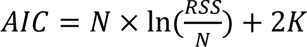

N is the length of the data, RSS is the squared sum of the residuals, and *K* is the number of free parameters used in the model fitting. We calculated the AIC for each model. When comparing the models, the model with the lower AIC that explained the data better was chosen. Additionally, statistical tests were performed with AIC values.

### Predicted Gain Test 1,2 and Multi-Predicted Gain Test

Predicted Gain Test 1,2 and Multi-Predicted Gain test have the following sequence of events that induce predicted gain (Fig. 2f,n,3a). Based on each event for each test, the parameters of each model are set to fit each model. We implemented the need model using the following equation.

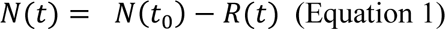

We assumed that the activity of neurons encoding need decreases in a stepwise manner as the animal goes through a series of events (Equation 17,18). We coded the need-encoding neurons with the structure of Equation 1. *N*(*t*) is the value for need at the time *t* and *N*(*t*_0_) is the value of need at the beginning of the experiment. *R*(*t*) indicates the stepwise increase of predicted gain, signaled by experimental events. *R*(*t*)is positive in these gain tasks.

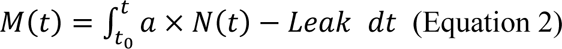

As the animal goes through a series of events, we assumed that the activity of the neurons that code for motivation is the integral of the state of need from the initial time to that point multiplied by the parameter ‘a’ (Equation 19) minus the value associated with ‘*Leak*’. Since ‘a’ and ‘*Leak*′ are fixed as constant for each event, and the value of need is also fixed at a constant value, the neurons that code for motivation can be coded as a linear function over each interval. We coded motivation with the structure of Equation 2.

#### a. Comparison with the model for individual event models

The equations used in the models are the same as the equations above, but the change in *R*(*t*) varies only once in the individual event model.

#### b. Multi-predicted gain/loss model

In the multi-predicted gain model, we additionally analysed changes in need and motivational state after the mice had finished eating (Fig. 4 and Extended Data Fig. 5). In this case, due to variability in the timing of food consumption in each trial, averaging by condition was not possible and individual trial analyses were performed (Extended Data Fig. 5). The individual trial analyses were performed with the same set of parameters, but computed RMSE across trials as mentioned above. The prediction of each model during the event that induced predicted gain/loss was the same as the previous equations. In the event that induced predicted loss, *R*(*t*) was a negative value (Equation 1,2).

### Predicted Loss Test 1

Predicted Loss Test 1 proceeded as follows: (Fig. 3a) In this test, only the event that induced predicted loss (inaccessible) exists. Based on the equations described above (Equation 1, 2), we implemented the code.

### Predicted Loss Test 2,3

Predicted Loss Test 2 has the following sequence of events (Fig. 3i). For each event, we set the parameters to match each model. Unlike Predicted Gain Tests, in Predicted Loss Test 2,3 the prediction was constrained to return to the baseline value before the trial start when the event that induced predicted loss was given (eating abandon). In this event, R(t) was 0. The equation of each model was described above (Equation 1,2).

### Behaviour Activity Model Preprocessing of Data

All data were analysed using MATLAB 2022B.

A probability of behavioural activity was calculated from periodic mouse behaviour annotations, by dividing a 1-second time period into bins: 10 bins for the eating-evoking test (Fig. 5). Whenever a behaviour occurred during a bin, the bin was recorded as “1.” Subsequently, all-time bins in a test per animal were averaged and normalised to derive a probability of behavioural activity. For each time bin, the behavioural activity was calculated as a moving average (10 bins for the 10 s stimulus test and 60 bins for the other bins) to create a smooth curve for model comparison.

### Model-Fitting Procedure

The need model defined that upon neuron activation at time *t*_*s* *on*_, the need state exhibited a value higher than its initial value. This higher need was continuously integrated to calculate the motivation level (*BN*(*t*), which was normalised according to the maximum value of the predicted data. *A* is the slope created when the stimulus-induced increase in need is integrated over time until time *t*_*s* *off*_. The motivation level was also calculated as a leaky integrated model based on the decrease in the need during the stimulus offset, such that the maximum decrease rate was not above 1/4^th^ of the increase rate during the stimulus on.

This calculated motivation level generated a behavioural activity when crossing the threshold (generated behaviour threshold). In addition, for the generated behaviour, the probability of the behaviour remained at a value of 1 because the motivation level continued to maintain the behaviour above a certain threshold (maintained behaviour threshold). Subsequently, the motivation level gradually decreased when the stimulus disappeared and only the effect of the intact *Leak* remained. Additionally, the probability of action decreased when the motivation level fell below the threshold (maintained behaviour threshold).

For the motivation model, the motivation level (*BM*(*t*) was observed to sharply rise (*A*_*stim*_) at the stimulus onset (*t*_*s* *on*_). The motivation level remained constant until the stimulus offset (*t*_*s* *off*_), after which it returned to the initial level.

Additionally, the code was implemented to compute the delay (*t*_*delay*_) between the actual neural activity and the behavioural results induced by the optogenetic stimulus and the threshold (generated behaviour threshold) using fmincon function in MATLAB for the best fit parameters with the smallest RMSE (Equation 3,4). Since the probability of an animal’s behaviour was modeled, the boundary values were set from 0 to 1. In the behavioural probability model, the threshold (maintained behaviour threshold) was set to 1, which meant that values greater than 1 were fixed at 1. In the model, the behavioural probability value was 1, but the state related to motivation was constantly increasing when the stimulus was on, and slowly decreasing when the stimulus was off due to a *Leak*. Similar to the process of the neural activity model, AICs were obtained from the predicted behaviour model and actual behaviour, and a statistical test was performed using the obtained AICs.

#### 1) Need

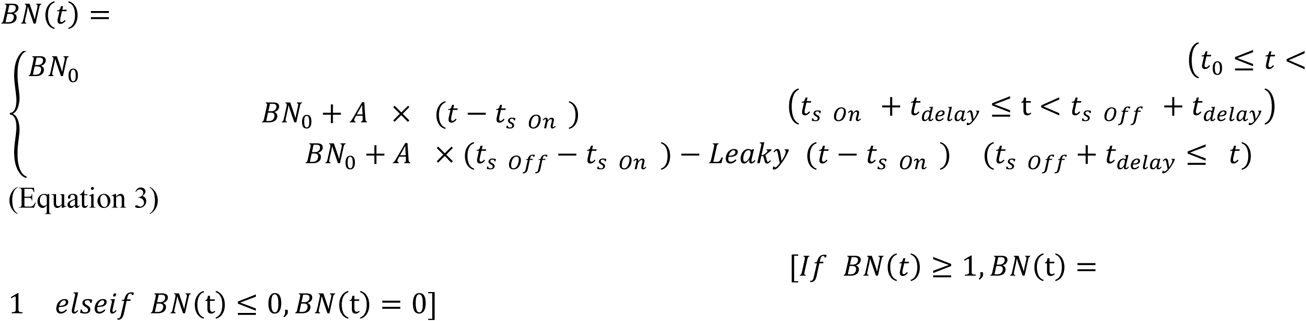

#### 2) Motivation

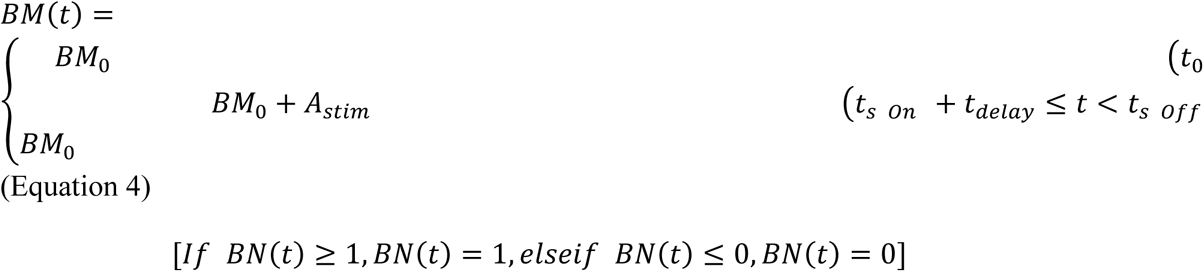

### Simulation Procedure

We examined different strategies of an agent using a computer simulation when the dynamics of need follows our normative framework. In the simulated environment, the agent forages for food resources over a long distance, experiencing a progressive increase in the deficit over time. This ensures the go/no-go decision-making process during foraging is induced by ecological factors that align with natural conditions. We tested the necessity of accumulation using two different simulation conditions.

#### 1) Need Model (Direct Need)

Actions were generated when the need value surpassed a threshold, and actions ceased when the need fell below the threshold.

1. Reward prediction changes when the agent stops In this situation, the agent can predict a decreasing deficit in the future while it is moving. However, if the agent stops, the agent immediately updates the prediction based on the increased future deficit.
2. Reward prediction is not affected by the motion of the agent In this situation, even if the agent stops, it computes a decreasing deficit due to the reward it will eat later.

#### 2) Motivation Model (Accumulated Need)

Actions were initiated when the motivation, the accumulation of need, exceeded a threshold. Actions were halted when the motivation value fell below the threshold.

1. lways accessible state condition The agent starts with the accessible condition. In these conditions the initial deficit at time 0 was assigned a value of 1 with subsequent increments of 0.00001 per time step.
2. With inaccessible state condition The agent starts in an initial state where the deficit remains at zero for 100 timesteps, after which an inaccessible state is introduced, causing the deficit to gradually increase over the next 500 timesteps. During this period of inaccessibility, the deficit increases linearly until it reaches a value of 1. Furthermore, in this condition, we conducted two simulations based on whether the deficit increases in the accessible state. This condition closely mimics the actual experimental setup, without introducing any additional events beyond accessible events in the experimental environment. In contrast to other simulations, we track the agent for 1,000 timesteps, starting from the moment the food is consumed, to illustrate the post-food consumption process. The deficit starts to decrease when the food is eaten by 0.012 per timestep and never falls below zero.

a. Increasing deficit during accessible condition (ID), increase the agent’s deficit by 0.00001 in the accessible state.
b. No increasing deficit during accessible condition (NID), during the accessible state, the deficit remains constant except for random noise.

#### 3) Simulation settings

1. The basis of simulation: The agent was modeled based on model rationality (Equation 1∼15).
2. Model compute: For the computation of SDD, r was defined as 0.99 (Equation 3). For the computation of PE, the parameter ‘e’ was specified as 0.1 (Equation 4), while the accumulation parameter ‘a’ for calculating motivation was set as 0.0015 (Equation 14). As motivation is calculated through the accumulation of need, it is set to decrease by 0.001 per 1 timestep for *Leak*. In the NID condition, *Leak* parameter was set 0.0008. For need value, the process was to treat values below 0 as 0.
3. Action generation: The threshold for generating the behaviour was defined as 0.5. We also considered the delay due to the behavioural switch between stay and go actions (*53*). The agent was set to maintain the stop behaviour at least 50 times steps after the switch. And for the go behaviour, it’s set to hold for 10 times steps.

### Statistical Tests

Statistical tests were performed by Friedman’s test with each model’s AIC value. A permutation test was performed if statistical significance was not secured in Friedman’s test due to insufficient data. The raw data was divided into 20 parts and each part was sampled without replacement to generate 1,000 sampling data of the same size as the raw data (Extended Data Fig. 4e–g). Subsequently, the model-fitting process was performed on the sampled data to obtain the AIC distribution of each model. Furthermore, this AIC distribution and the AIC of the actual data were used for obtaining the empirical p-value (Extended Data Fig. 4f, g).

### Model Permutation Test Procedure

To verify our model trace fit with experimental data rather than random data, raw calcium traces or behaviour activity were shuffled into a random data set. Afterward, the residual sum of squares was calculated for the model-random data set and the model-experimental data set. Afterward, the permutation test was permuted 1,000 times between the two data sets.

### PCA Analysis

For preprocessing, neural activity was collected from the multi-predicted gain test trials from AgRP or LH^LepR^ neurons. All the trials were subtracted with its corresponding average value from the whole trial and concatenated to a matrix (3,000 frames per trial) to be processed with MATLAB function PCA. The product of principal component and individual neural activity (coeff) was used to make the neural trajectory for analysis. PC1, PC2, PC3 explained about 80.9 %, 9.2 %, 3.6 % of the variance, respectively. Afterwards, PC1 data was processed with MATLAB’s function tsne with perplexity 10 for t-SNE distribution results.

### CEBRA Analysis

For preprocessing, neural activity was collected from the multi-predicted gain test trials from AgRP or LH^LepR^ neurons. The trials were reshaped to a matrix form comprising three trials (750 frames per trial) × n dimension (n for AgRP = 24, n for LH^LepR^ = 19). Afterward, the neural activity was up-sampled to 100 Hz per second (Total 3,000 frames). Behaviour label was differentiated to continuous label scores for all-time frames to match the number of columns of the reshaped trial matrix: trial start = 0 (0s) /accessibility moment = 1/seeking moment = 2/proximate to food = 3/contact = 4/trial end = 5 (the 30s). After preprocessing, 70% of the trial matrix and continuous behaviour labels were split for the training set and the rest were designated test sets. Continuous behaviour labels were then shuffled to make shuffled sets for model fitting. The preprocessed data sets were used in the CEBRA modeling function *(46)* that was built with the following parameters: learning rate: 0.00005, maximum iteration: 100,000, temperature_mode: auto, model_architecture: “offset10_model,” batch_size: 1024, outpud_dimension = 3, time_offsets = 10, conditional = “time_delta.” The plot of temperature change and loss was used with functions cebra.plot_loss and cebra.plot_temperature. Consistency scores between hypothalamic neurons were computed using the consistency_score function. Behaviour prediction was decoded from embeddings using, k-nearest-neighbour (kNN) decoder. The prediction was used with real labels to compute RMSE for comparison between original and shuffled data.

## Theoretical Backgrounds

In the normative approach of homeostatic theory in all homeostatic contexts, individual contexts constitute independent axes in a multidimensional space H. An animal’s deficit (*D*_*i*_(*H*_*t*_)) in homeostatic axis *i* and time *t*, is defined as the degree to which a state deviates from a homeostatic setpoint 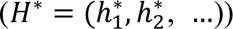 and homeostatic state(*H*_*t*_ = (ℎ_1,*t*_, ℎ_2,*t*_, …)) (*7*). Here, we redefine *D*_*i*_(*H*_*t*_) as below (Equation 5). The newly defined *D*_*i*_(*H*_*t*_) takes the difference between setpoint and state from each axis that affects homeostasis, and considers the impact of each factor (e.g., nonlinearities) independently through a function *f*_*i*_. Then, the total deficit at time *t* (*D*_*total*_(*H*_*t*_)) is determined as the linear sum of the deficit from each axis (Equation 6).

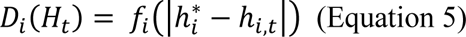

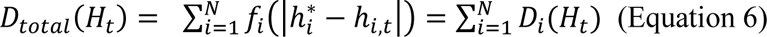

An internal state deviating from the homeostatic set point is perceived as a current deficit (*8*). The animal analyses external information to predict future events, thereby forecasting its future predicted deficit in addition to its current deficit (*9*–*14*) The core task of the brain is to anticipate an animal’s future deficits, regulating internal state for preparation of deficit changes. Therefore, for optimal survival, the predicted deficit should serve as the optimal update for change in state. The cumulative effect of *D*_*total*_(*H*_*t*_) over time, as the state of homeostasis progresses along the trajectory (policy π), is defined as the sum of the discounted deficit, and can also be understood as the sum of all discounted deficit (*SDD*_π_(*H*_*t*_)) (Equation 7).

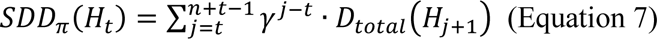

*SDD*_π_(*H*_*t*_) is a value that predicts information about a future state based only on what is happening at the time *t*.

If the animal is given accessibility at time *t*_*acc*_, the animal will try to minimise its deficit, and this process will cause a change in an animal’s prediction (Fig. 1a). We define this amount as predicted change (*PC*_*total*_(*H*_*t*_)) (Equation 3). The predicted change at time *t* can be found by multiplying the difference between the sum of all discounted deficit at time *t* and time *t*_*acc*_by a free parameter, *w*. The total predicted deficit (*PD*_*total*_(*H*_*t*_)) can be described as the sum of the total predicted deficit at the current state and the total predicted change (Fig. 1) (Equation 9).

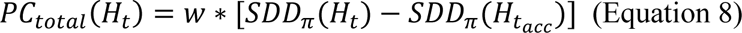

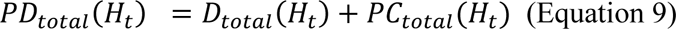

According to the normative approach in homeostatic theory, which integrates homeostasis theory and reinforcement learning, the deficit during each state transition within homeostasis serves as a reward for reinforcement learning (*7*). This reward can be expressed as the sum of discounted rewards (*SDR*_π_(*H*_*t*_)) by the state-value function (Equation 10). If the behavioural policy that maximises *SDR*_π_(*H*_*t*_) aligns with the policy that minimises *SDD*_π_(*H*_*t*_), the animal can maintain homeostasis and learn through reinforcement, which is optimal for efficient survival (Equation 11,12).

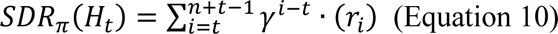

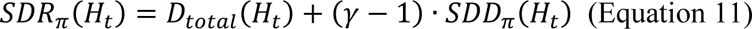

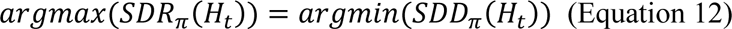

Since, according to our new definition of deficit (Equation 5,6), *SDD*_π_(*H*_*t*_) and *PD*_*total*_(*H*_*t*_) are the linear sum of the differences along each homeostatic axis, the *PD*_*total*_(*H*_*t*_) can also be expressed by linearity as the sum of the *PD*_*i*_(*H*_*t*_) along each axis. When there are N homeostatic axes, the *PD*_*total*_(*H*_*t*_) can be expressed as (Equation 13).

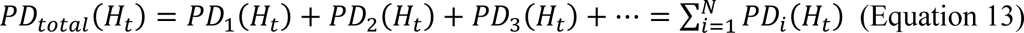

Minimising the total predicted deficiency, *PD*_*total*_(*H*_*t*_), would be important for the animal’s survival. If a change is induced in only one axis (e.g., axis 1), such as in our controlled experiments, the policy that minimises the *PD*_1_(*H*_*t*_) on this one axis will be equivalent to the policy that minimises the *PD*_*total*_(*H*_*t*_). Also, these policies correspond to the ones that maximise the *SDR*_π_(*H*_*t*_) (Equation 14).

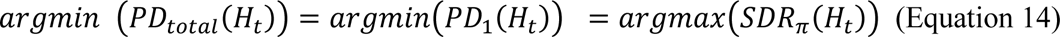

Also, in controlled experiments with minimal food intake, the current deficit state has the same value at each time point as the trial progresses.

Note that the above relationship still holds in the previous study (*7*) in certain conditions. In the previous study, *D*(*H*_*t*_) was defined as follows:

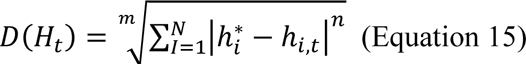

When the free parameter m (Equation 11), which determines the *D*(*H*_*t*_), is set to 1, *D*(*H*_*t*_) can be expressed as the linear sum of homeostatic *D*(*H*_*t*_) on each axis of the factors influencing homeostasis. In this case, the *PD*_*total*_(*H*_*t*_) can be divided by the *PD*_*i*_(*H*_*t*_) of each homeostatic factor. Furthermore, in our controlled experiments, the homeostatic *D*(*H*_*t*_) may be mainly along a single context axis (i.e., food) that affects homeostasis (m=n=1; Equation 15). In such cases, *PD* can be considered as a factor that solely responds to variations of this homeostatic axis (food).

Within the realm of feeding, the current “food deficit” is *D*_*f*_(*H*_*t*_) (Equation 5, Extended Data Fig. 1b Phase 2), while the changes in the sum of discounted future “food deficit” represents the sum of predicted changes (gain or loss) *PC*_*f*_ that alarm feedforward signals (Equation 8, Extended Data Fig. 1b Phase 3) to sum up to a predicted deficit for food (*PD*_*f*_(*H*_*t*_)) (Equation 17).

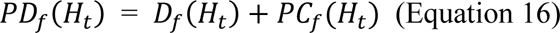

It is known that the bodily system of an animal responds to the cue from the environment which might lead to homeostasis recovery to produce a predicted body state *PD*_*f*_(*H*_*t*_) (*54*–*56*). This anticipatory regulation system is paramount for survival because it generates real-time feedforward signals that prevents overcompensation of the bodily system when the bodily system changes back to the set point after state deviation (*15*, *54*, *57*). Consequently, in the hunger domain, *PD*_*f*_(*H*_*t*_) should serve as the animal’s optimal calculation for food “need” *N*_*f*_(*t*) (Equation 17).

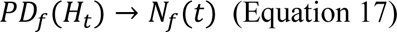

In the domain of animal behaviour and survival, animals constantly decide whether to seek for food or not, by combining complicated external cues and interoceptive states that are often uncertain. Animals often produce behaviours through accumulation of these cues (*3*, *28*, *29*). Also, recent research has shown that cognitive features such as needing and wanting could be separately dissociated in computational simulation (*40*). To explore the question of whether accumulation is necessary under the constraints of our normative framework, we conducted a computer simulation to discern whether need requires accumulation for optimal eating behaviour.

In an experiment based on need, two rules were put to the test. The first rule suggested that the predicted reward for the agent should change based on its behavioural states, while the second proposed that the predicted reward should not be affected by the state of behaviour.

Following the first rule, the agent began its search for food from a starting point. As it got close to the food, the expected reward increased, causing the agent’s need to decrease. Once the need fell below a threshold, the agent stopped and triggered a reset in reward expectation. This reset caused the need to rise rapidly again, albeit with a delay due to a behavioural change. As the agent resumed movement, the agent’s internal variables reset again, causing the need to decrease even more rapidly and to a lower level than before. This resulted in a cycle of starts and stops, ultimately making the agent largely inefficient in the food-seeking behaviour (Supplementary Movie 1, Extended Data Fig. 1b).

The results of the second rule were similar to those of the first, as the agent’s initial behaviour was the same. However, unlike the first rule, the system did not reset when interrupted by a decline in need. Consequently, the agent’s deficit gradually increased, leading to a slow increase in need. Eventually, when the need surpassed the threshold, the agent started moving again. However, as soon as the agent moved, the need fell below the threshold, trapping it in a prolonged loop, similar to the first rule (Supplementary Movie 1, Extended Data Fig. 1c). While repeatedly performing go-and-stop actions may eventually lead to a reward, this alternating behaviour can be inefficient and even dangerous. Energy may be wasted, and the agent may be exposed to predators, putting its survival at risk.

On the other hand, actions driven by motivation had a slower start by accumulating need. However, it exhibited strong persistence once motivation reached a threshold. Even though the need decreases (or even goes to zero) as the agent gets close to the food, the motivation remains above the threshold due to the accumulation of need. This unwavering motivation successfully guided the agent towards eating food, surpassing the performance of the need model (Supplementary Movie 1, Extended Data Fig. 1d,e). These simulation results underscore the advantage of motivation-driven behaviour over solely need-based behaviour.

Therefore, motivation should be optimally generated as the integration of need. Motivation is also influenced by an integrated affecting factor (*a*). For example, motivation does not accumulate before accessibility is granted (*24*) (*a* = 0). Additionally, motivation changes based on factors, such as satiety, valence, and memory (*24*, *58*) (*a* ≥ 0). Therefore, the food-directed motivation at a specific time point (*M*_*f*_(*t*)) can be expressed as the integral of the product between the integrated affecting factor and need. Finally, a leaky integrator was included to account for factors (mentioned above) that continuously devalue motivation in an animal.

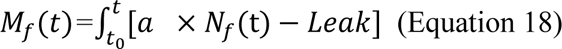

When sufficient motivation accumulates and surpasses a certain threshold (K ≥ 0), behaviour activity starts manifesting (*30*). Therefore, behaviour outcome at a certain time point (*t*), can be written as follows:

## Data and code availability

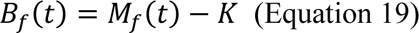

All data and code will be provided upon request to the corresponding author

## Acknowledgements

We thank all the lab members of FNMR Lab and Ben Dougen (NeuRLab) for technical assistance, valuable discussions and feedback on the manuscript.

## Author Contributions

K.S.K., Y.H.L., Y.B.K., J.W.Y., H.R.K. and H.J.C. designed the project. K.S.K., Y.H.L., Y.B.K., and J.W.Y wrote the draft and performed visualization of data. All other authors discussed the results and further edited the manuscript. Y.B.K., Y.H.L. and H.Y.S. performed stereotaxic surgeries and brain imaging. K.S.K., Y.H.L., Y.B.K., H.Y.S. and S.H.J. performed the behavioural test and behaviour labeling. J.W.Y. performed model-fit analysis with assistance from K.S.K.; K.S.K. and J.S.P. provided initial version. K.S.K. performed the unsupervised dimensionality reduction analysis. J.W.Y performed the computer simulation. K.S.K. and J.W.Y. performed the statistics. J.W.S. and K.W.K. provided mouse lines for the study. H.R.K. and H.J.C supervised the project.

## Additional Information

Supplementary Information is available for this paper.

**Extended Data Fig. 1.**
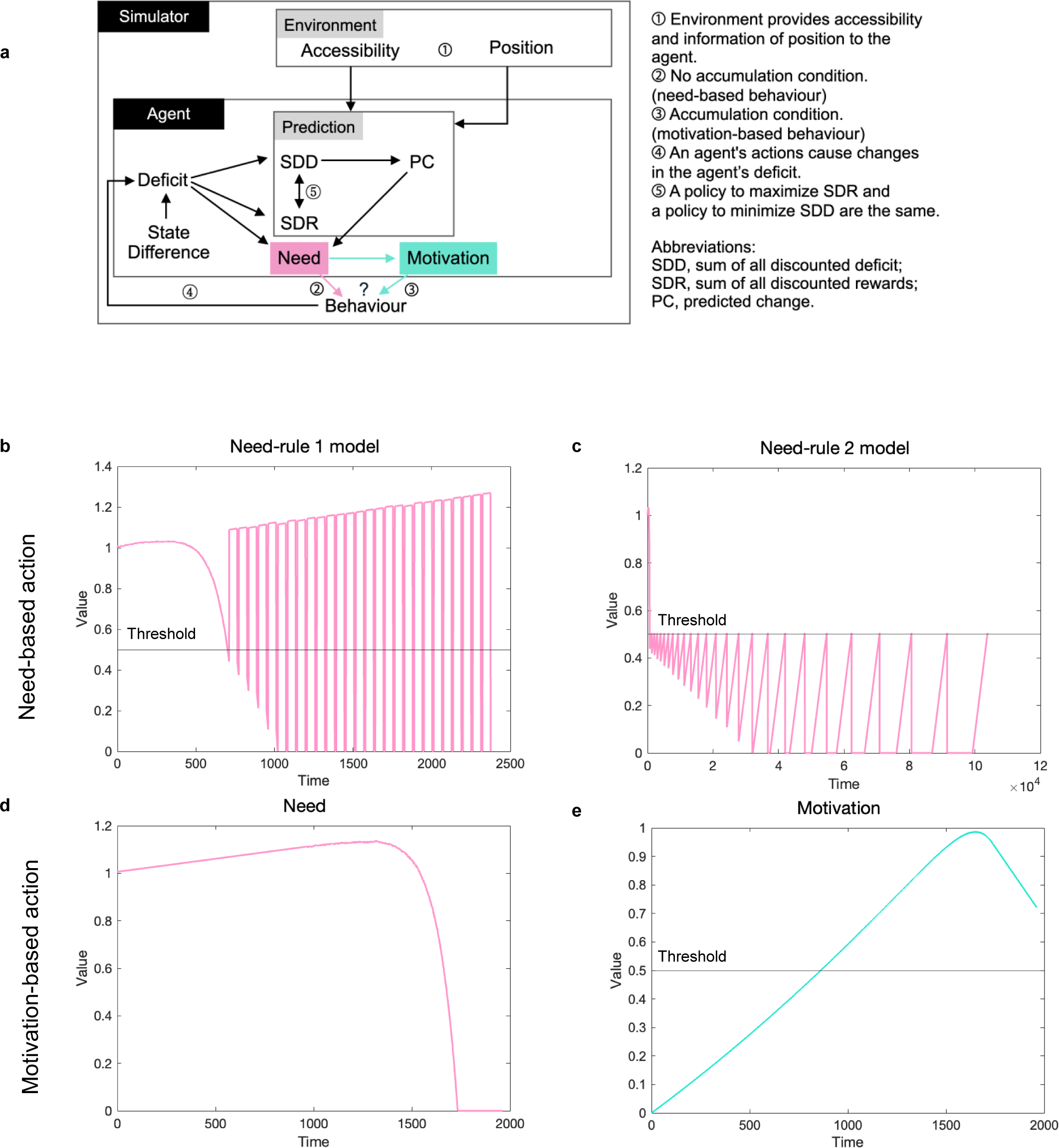
Simulations Demonstrate Accumulation of Need is Required for Survival. **a**, The schematic of the simulation. **b, c,** Time courses of the entire progression in the model where behaviour is generated by need. Both models exhibit an inefficient pattern of repeated go and stop based on the threshold. The black line is the threshold. **d, e,** Time courses for the entire progression in the model where behaviour is generated by motivation. The black line is the threshold.

**Extended Data Fig. 2.**
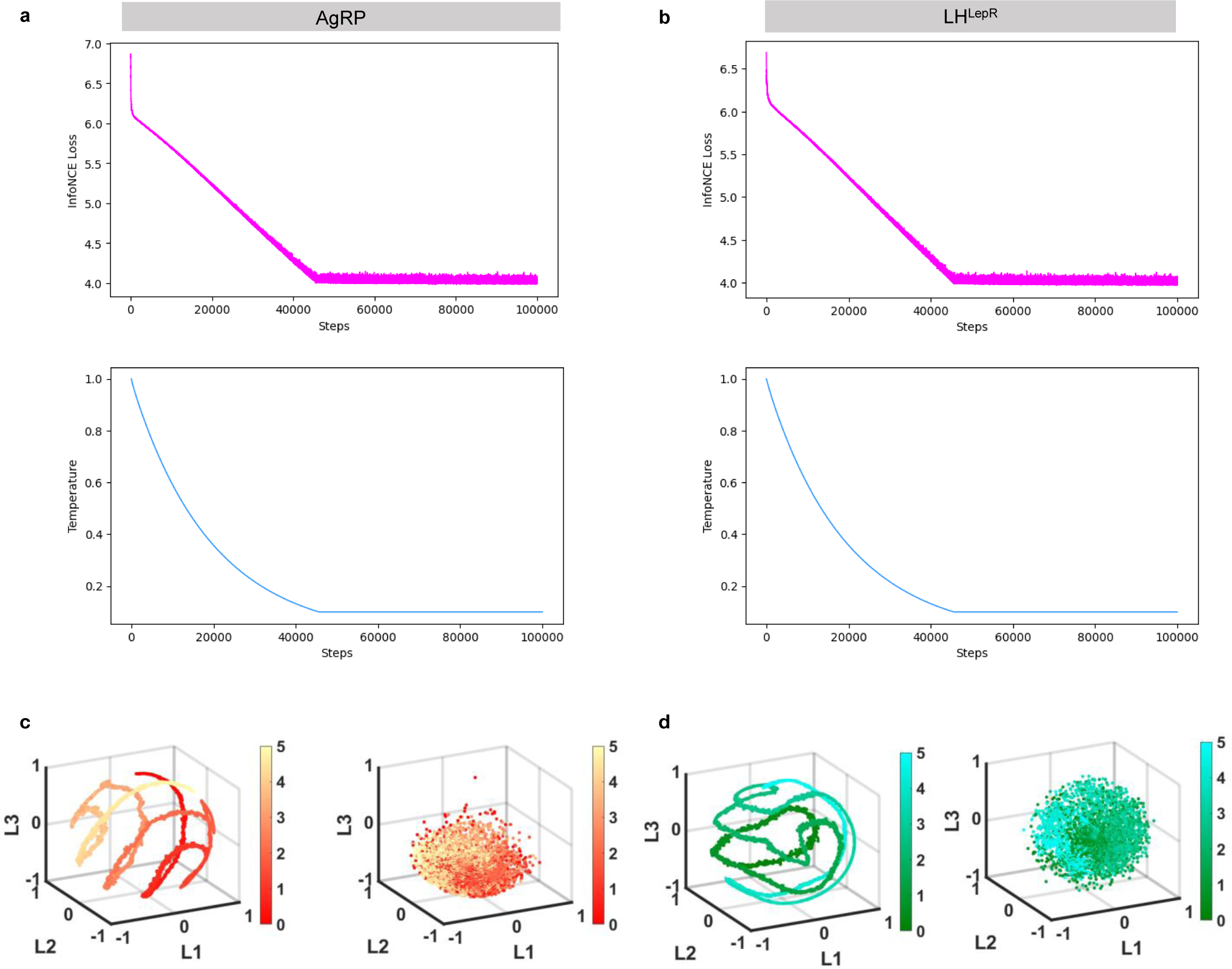
CEBRA Analysis Show Different Hidden Embeddings in Hypothalamic Neurons. **a, b**, Convergence of training loss and temperature for model evaluation. **a,** AgRP neurons (N = 6, Trials = 72), **b,** LH^LepR^ neurons (N = 4, Trials = 56). **c, d,** Embeddings from CEBRA analysis from hypothalamic neurons. Left are embeddings from original labels, right are embeddings from shuffled labels. Colourbar is shown from 0 to 5 in order of predicted gain events (See the Methods). **c,** Embeddings from AgRP neurons (N = 6, Trials = 72). **d,** Embeddings from LH^LepR^ neurons (N = 4, Trials = 56). See Supplementary Table. 1 for statistics.

**Extended Data Fig. 3.**
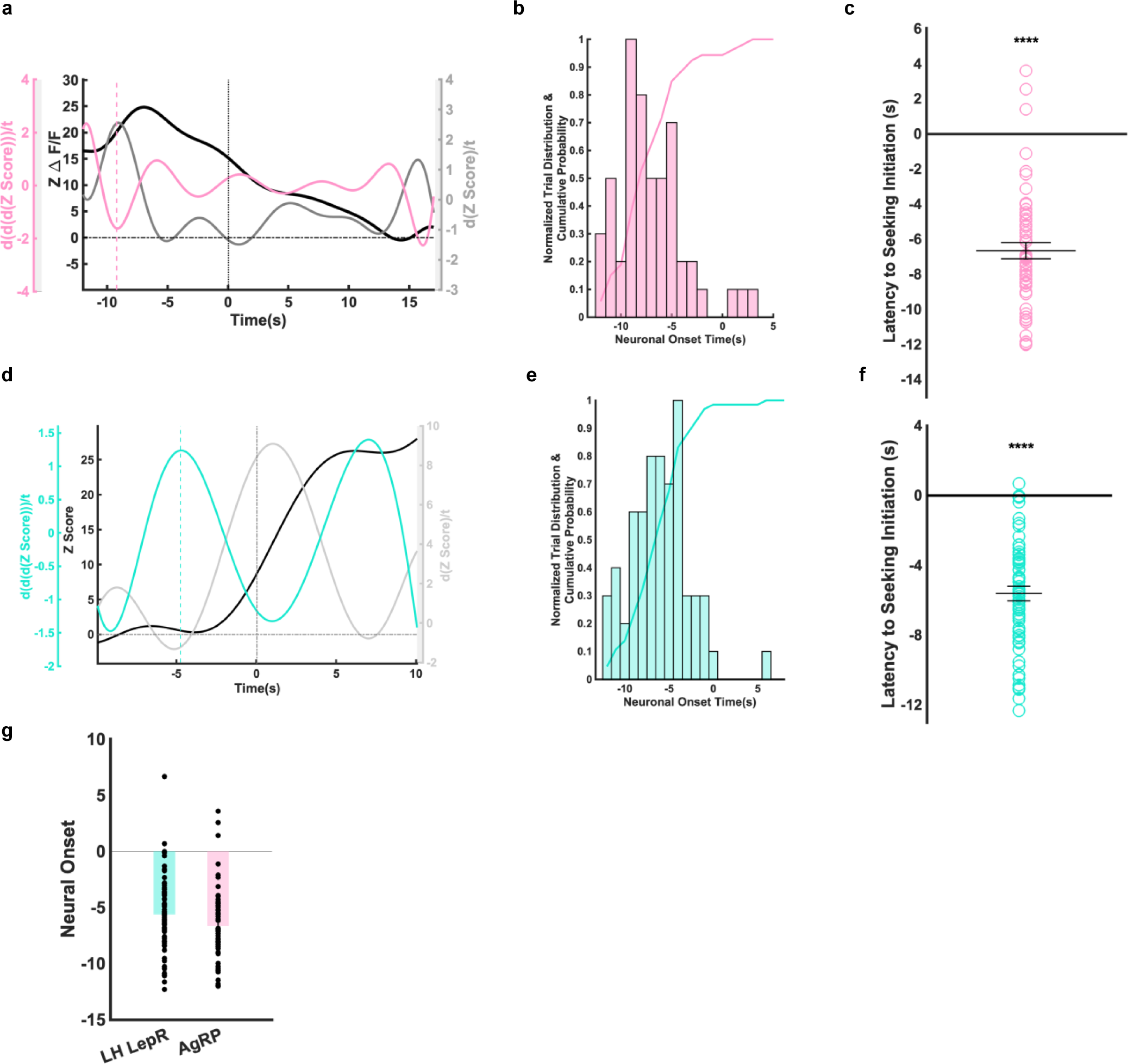
Neural Onset of AgRP and LH^LepR^ Neurons Show Temporal Sequence of Neural Activity. **a, d,** Representative graph of neural onset from AgRP or LH^LepR^ neurons from the predicted gain test 1. Black is neural activity. Grey is the first derivative and pink or turquoise is the third derivative of neural activity. **a,** Individual neural activity data from AgRP neurons, **d,** Individual neural activity data from LH^LepR^ neurons. **b, e,** Histogram and cumulative probability from AgRP or LH^LepR^ neural activity. **b,** AgRP neural activity (N = 4 Trials = 54), **e,** LH^LepR^ neural activity (N = 5 Trials = 65). **c, f,** Quantification of latency to seeking initiation after neural onset from AgRP or LH^LepR^ neurons. **c,** AgRP neurons, **f,** LH^LepR^ neurons. **g,** Comparison of neural onset between AgRP (pink) and LH^LepR^ (turquoise) neurons. Data are mean ± s.e.m. See Supplementary Table. 1 for statistics.

**Extended Data Fig. 4.**
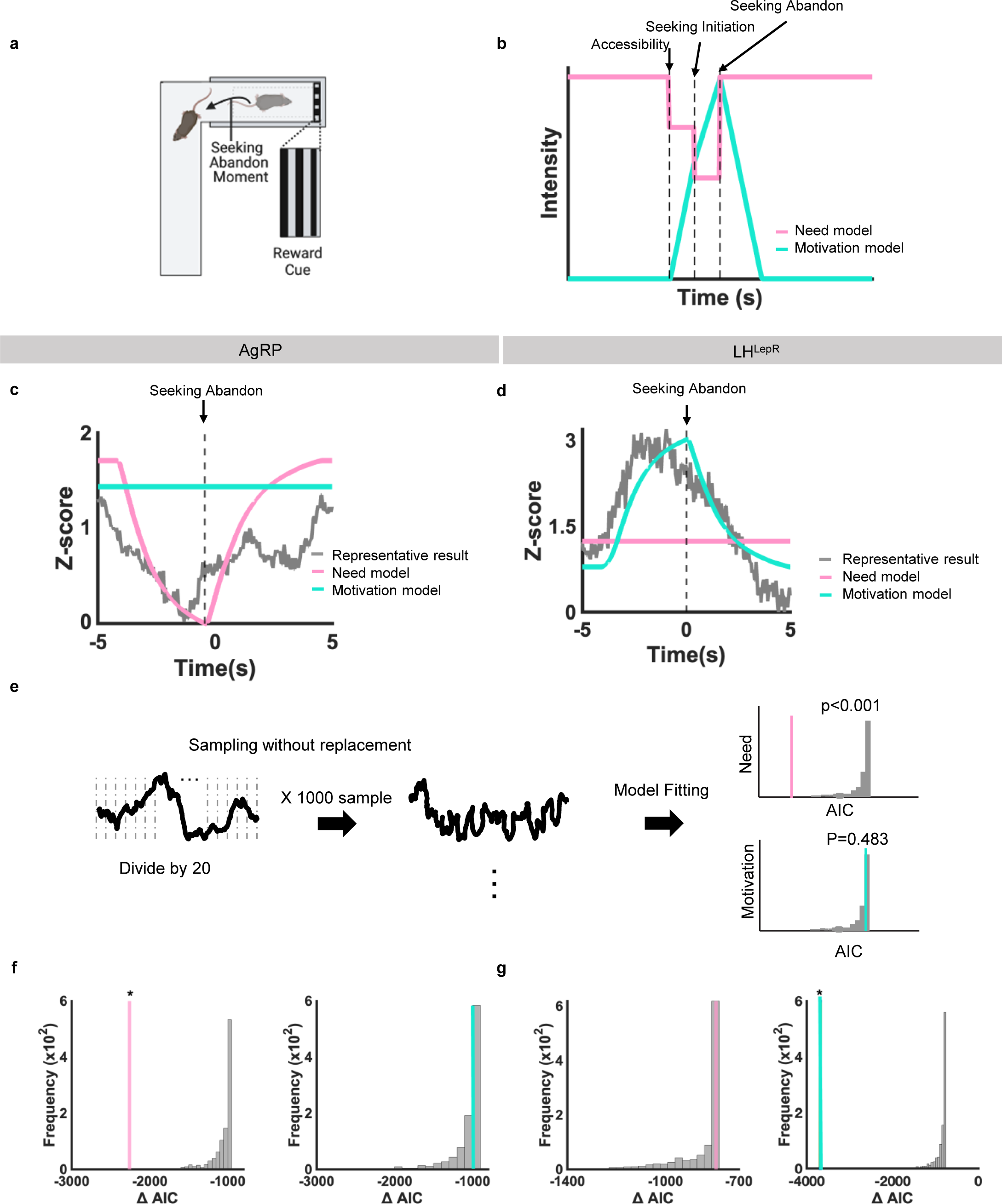
Individual Trial Model Fitting Results Dissociates Need and Motivation in the Hypothalamic Neurons. **a**, Schematic of behavioural paradigm and event moment. **b,** Schematic of model prediction of neural activity from need (pink) and motivation (turquoise) neurons. **c, d,** Individual trial fitting of neural activity. Need neural activity model is pink and motivation neural activity model is turquoise. Grey is neural activity (normalised Z-score). Dotted line indicate seeking abandon moment. **c,** Neural activity of AgRP neurons, **d,** Neural activity of LH^LepR^ neurons. **e,** Schematic of individual trial analysis. **f, g,** Permutation test result of model fitting. Histogram is AIC from difference between model and sampled data set or experimental data set. **f,** Result from AgRP neural activity (N = 1, Trials 5, permuted 1000 times), **g,** LH^LepR^ neural activity (N = 1, Trials 5, permuted 1000 times). See Supplementary Table. 1 for statistics. The schematics in **a** were created using BioRender.

**Extended Data Fig. 5.**
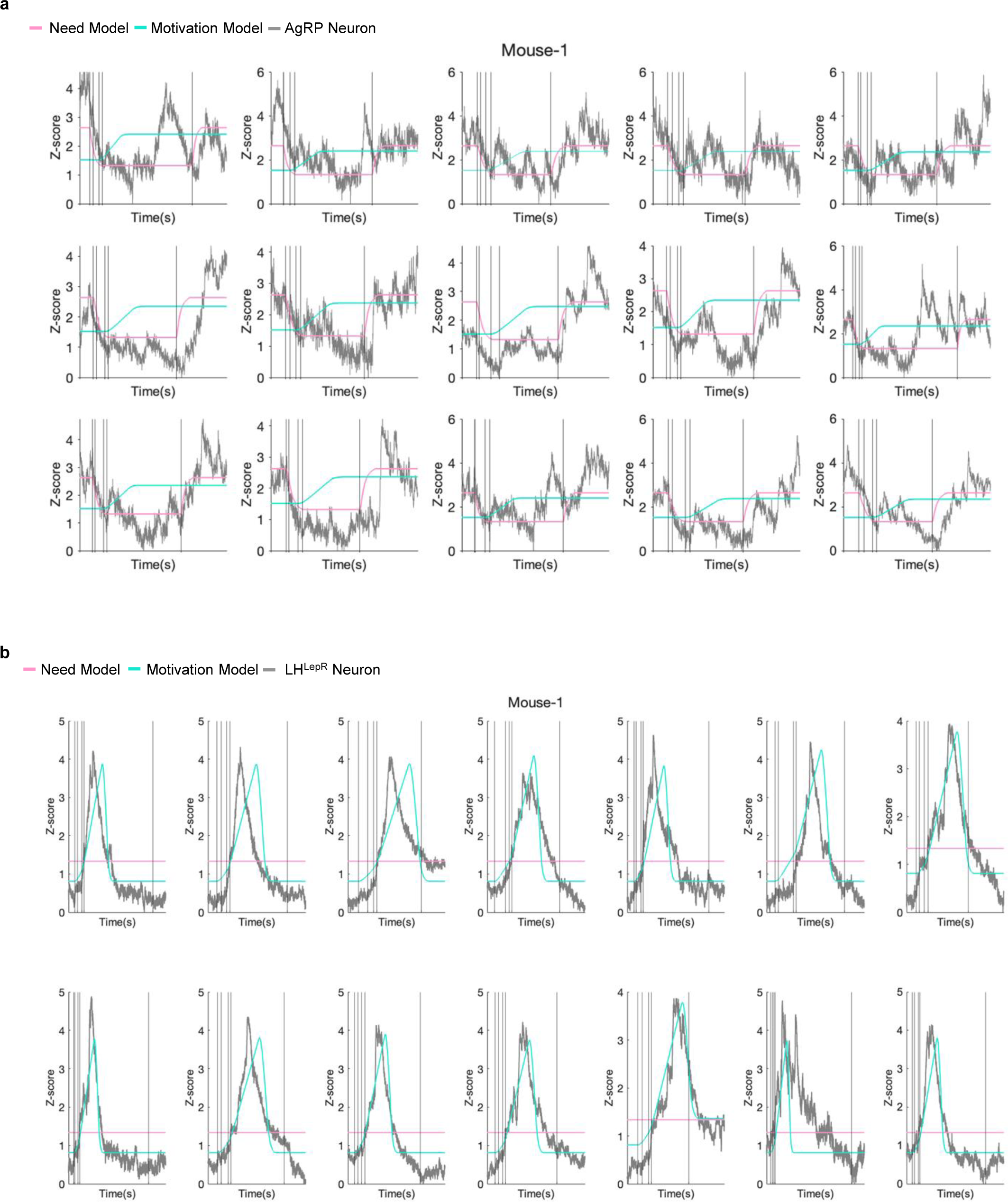
Model Fitting of Individual Trials from AgRP or LH^LepR^ Neural Activity Consistently Dissociate Need and Motivation, Respectively. **a, b,** Model fitting of individual trials from AgRP neurons. Grey line is neural activity (normalised Z-score), turquoise is optimal model fit for motivation neural activity, pink is optimal model fit for need neural activity. Vertical lines along x-axis correspond to accessibility, seeking initiation, proximate to food, contact, inaccessibility moment in order. **a,** AgRP neurons, **b,** LH^LepR^ neurons.

**Extended Data Fig. 6.**
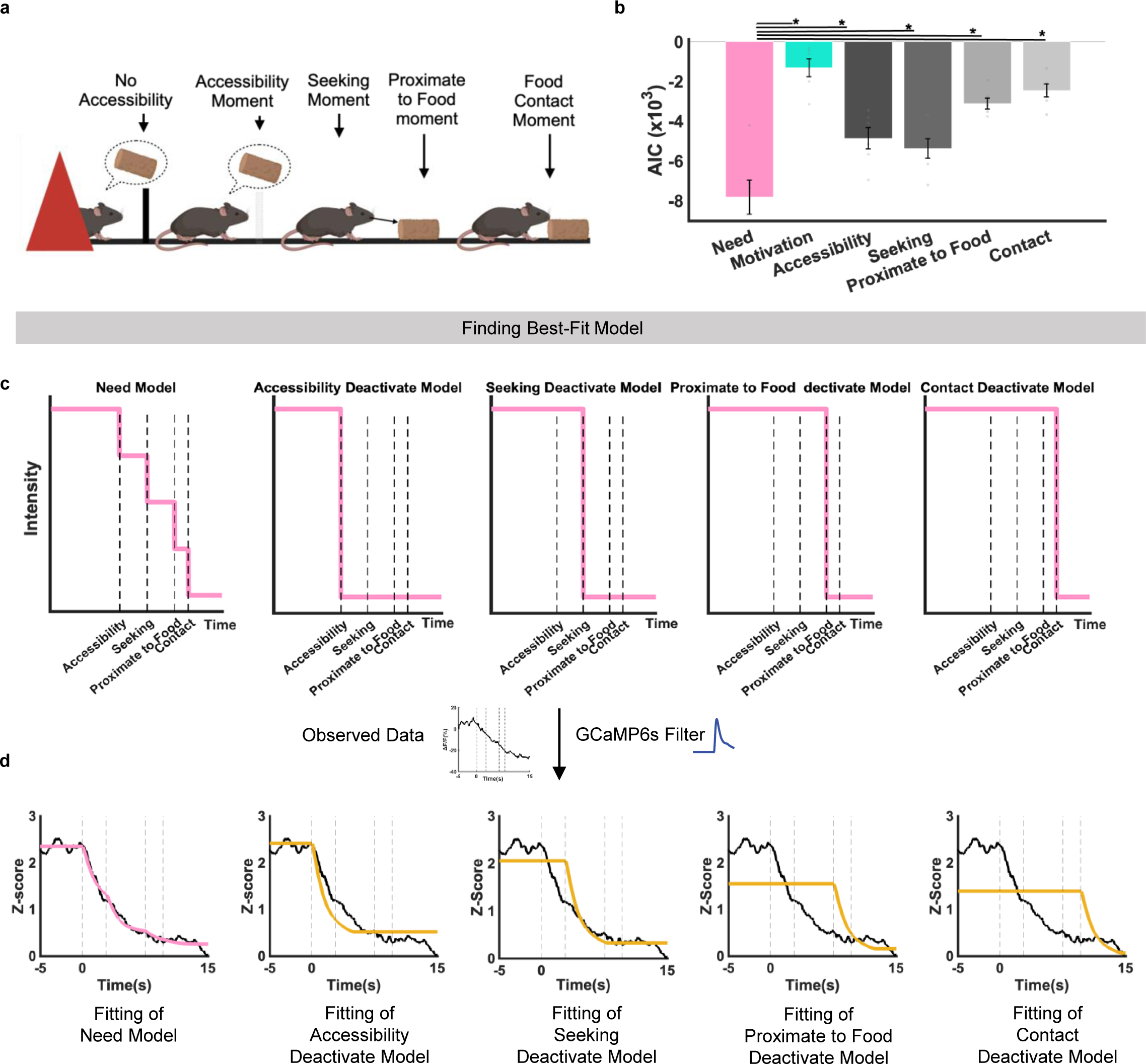
Model Fitting Between Candidate Models and AgRP Neural Activity Show Need Neural Activity Model as Best-Fit Model. **a**, Schematic of behavioural paradigm and event moment during multi-predicted gain test. **b,** Quantification of AIC between AgRP neural activity and candidate models (N = 6). **c,** Schematic of model prediction from each candidate model. **d,** Representative model fitting between average trace of AgRP neural activity (normalised Z-score) and candidate models from 1 mouse. Dotted line along x-axis correspond to accessibility, seeking, proximate to food, contact moment in order. Data are mean ± s.e.m. See Supplementary Table. 1 for statistics. The schematics in **a** were created using BioRender.

**Extended Data Fig. 7.**
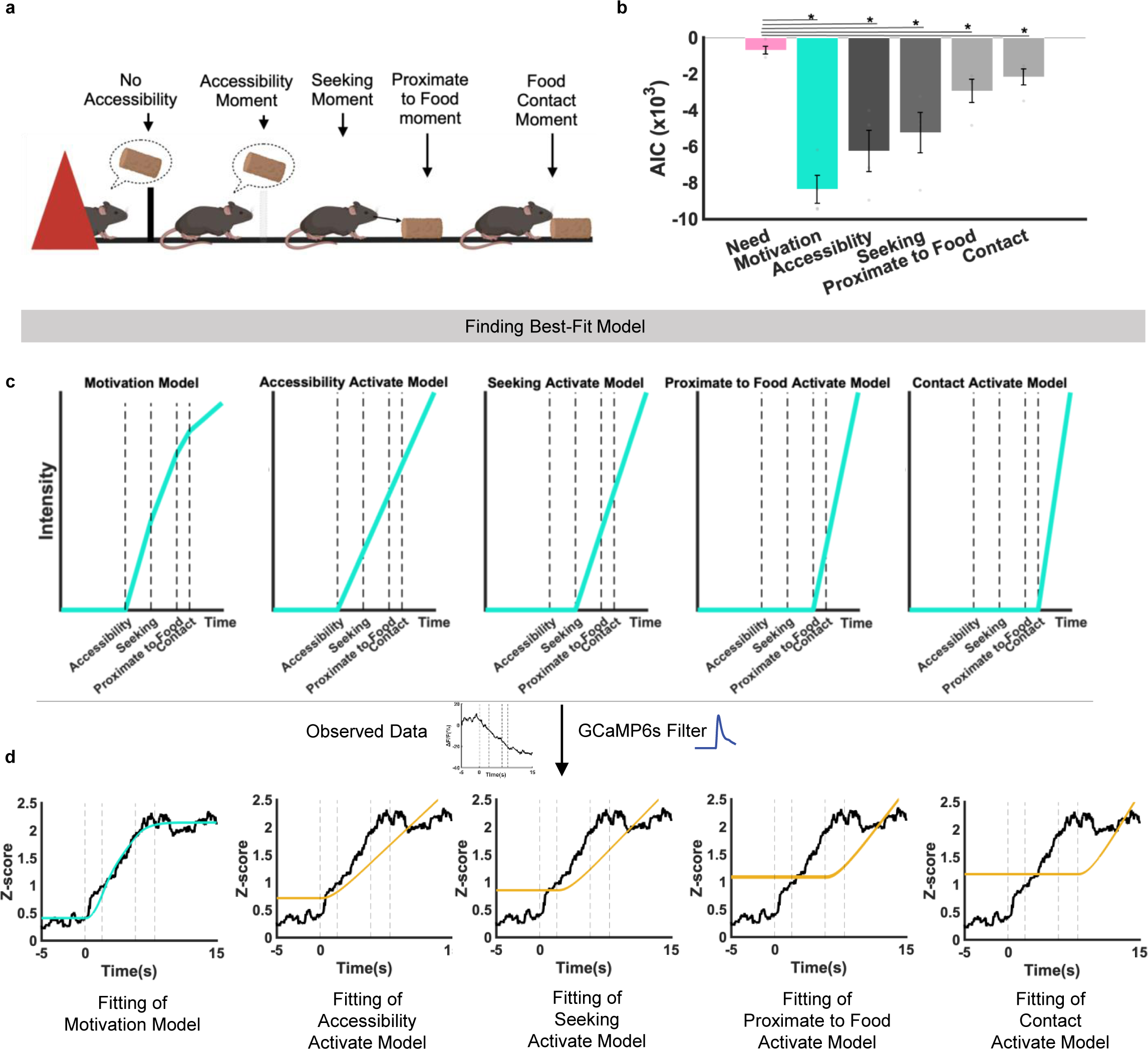
Model Fitting Between Candidate Models and LH^LepR^ Neural Activity Show Motivation Neural Activity Model as Best-Fit. **a**, Schematic of behavioural paradigm and event moment multi-predicted gain test. **b,** Quantification of AIC between LH^LepR^ neural activity and candidate models (N = 4). **c,** Schematic of model prediction from each candidate model. **d,** Representative model fitting between average trace of LH^LepR^ neural activity (normalised Z-score) and candidate models from 1 mouse. Dotted line along x-axis correspond to accessibility, seeking, proximate to food, contact moment in order. Data are mean ± s.e.m. See Supplementary Table. 1 for statistics. The schematics in **a** were created using BioRender.

**Extended Data Fig. 8.**
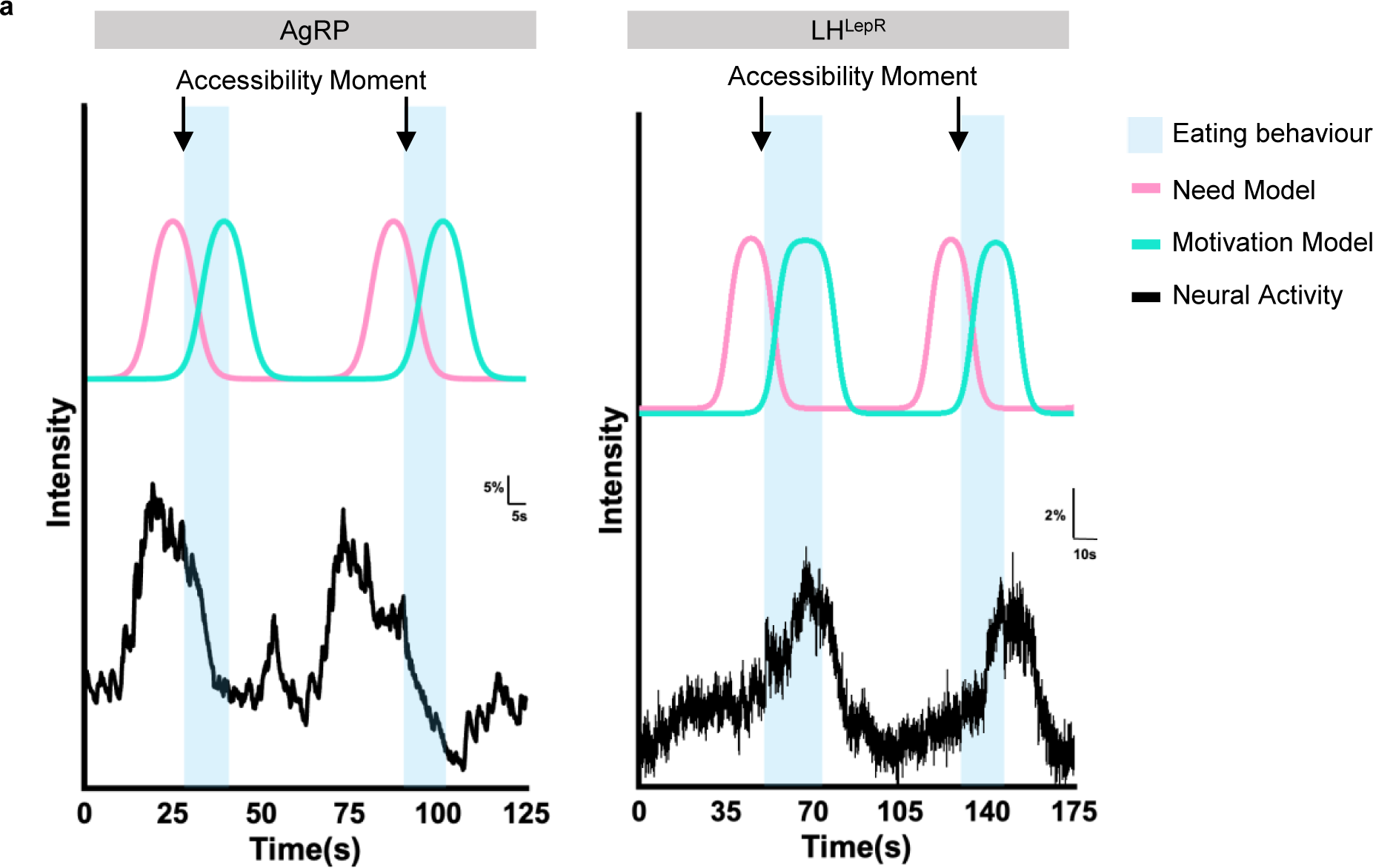
Summary of the Temporal Dynamics of AgRP and LH^LepR^ Neurons Encoding Need and Motivation, Respectively. **a**, Schematic of AgRP neurons (black, left) and LH^LepR^ neurons (black, right) encoding need (pink) and motivation (turquoise), respectively.

**Supplementary Table. 1.**
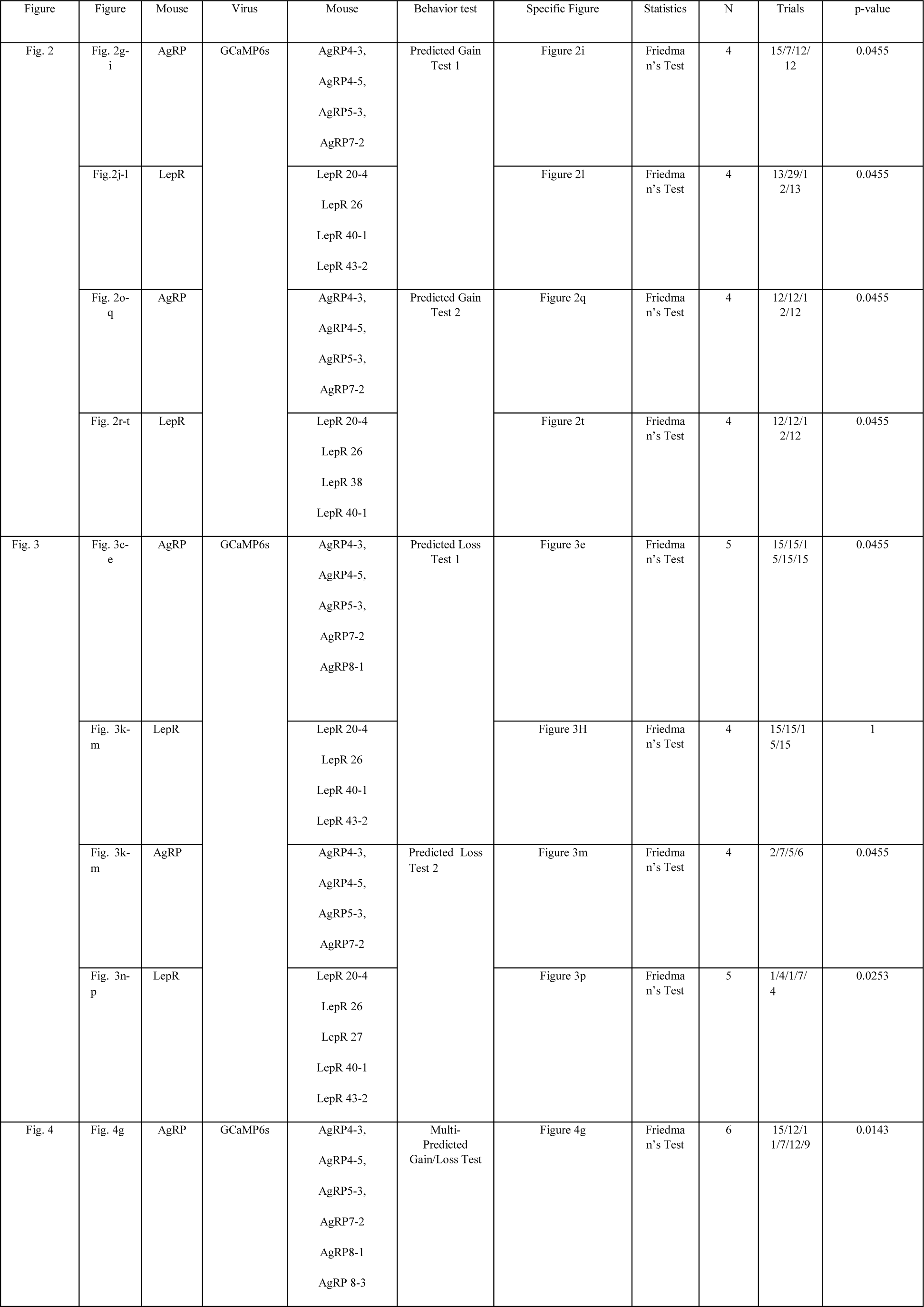
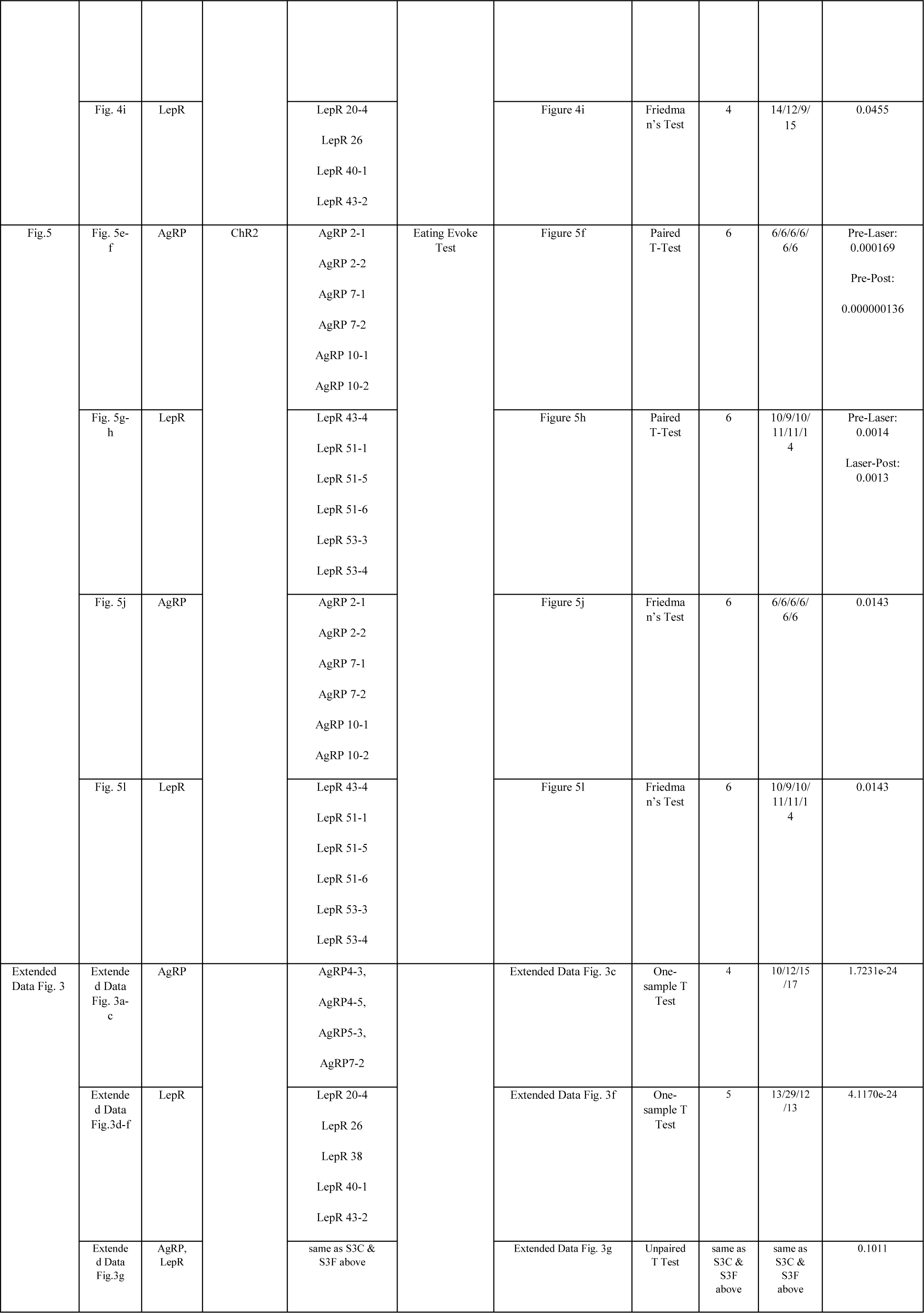
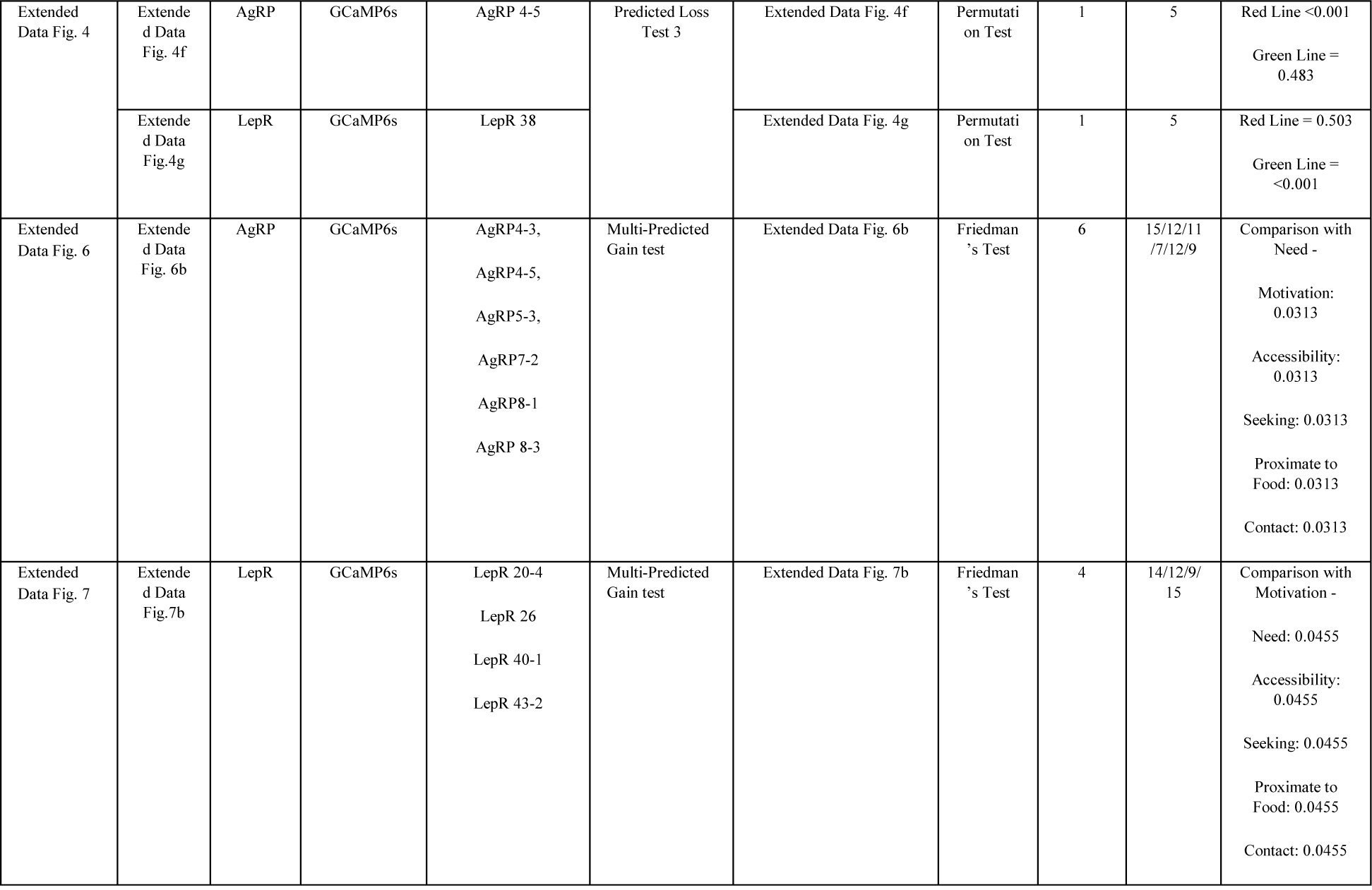
Summary of statistical analyses.

## Supplementary Video Legend 1 to 5

Supplementary Video 1. Selection of Need-Driven Behaviour Policy from Motivation-Driven Behavior Policy Using Computer Simulation

Supplementary Video 2. CEBRA Analysis Reveals Different Latent Features in AgRP and LH^LepR^ Neurons

Supplementary Video 3. Neural Activity Model Prediction of AgRP and LH^LepR^ Neurons Reveals Role as Need and Motivation, Respectively

Supplementary Video 4. Behaviour Activity Prediction of AgRP and LH^LepR^ Neurons Reveals Role as Need and Motivation, Respectively

Supplementary Video 5. Summary of Temporal Dissociation of the Role of Hypothalamic Neurons as Need and Motivation

## References

1. W. B. Cannon, The wisdom of the body. (W.W. Norton & Company, New York,, 1932), pp. xv p., 1 l., 19–312 p.

2. Y. H. Lee et al., Food Craving, Seeking, and Consumption Behaviours: Conceptual Phases and Assessment Methods Used in Animal and Human Studies. J Obes Metab Syndr 28, 148–157 (2019).

3. G. Pezzulo, F. Rigoli, K. Friston, Active Inference, homeostatic regulation and adaptive behavioural control. Prog Neurobiol 134, 17–35 (2015).

4. W. E. Allen et al., Thirst-associated preoptic neurons encode an aversive motivational drive. Science 357, 1149–1155 (2017).

5. C. L. Hull, Principles of behaviour: an introduction to behaviour theory. Principles of behaviour: an introduction to behaviour theory. (Appleton-Century, Oxford, England, 1943), pp. x, 422-x, 422.

6. K. Juechems, J. Balaguer, S. Herce Castanon, M. Ruz, J. X. O’Reilly, C. Summerfield, A Network for Computing Value Equilibrium in the Human Medial Prefrontal Cortex. Neuron 101, 977–987 e973 (2019).

7. M. Keramati, B. Gutkin, Homeostatic reinforcement learning for integrating reward collection and physiological stability. Elife 3, (2014).

8. K. C. Berridge, Motivation concepts in behavioural neuroscience. Physiol Behav 81, 179–209 (2004).

9. C. A. Zimmerman et al., Thirst neurons anticipate the homeostatic consequences of eating and drinking. Nature 537, 680–684 (2016).

10. D. S. Ramsay, S. C. Woods, Clarifying the roles of homeostasis and allostasis in physiological regulation. Psychol Rev 121, 225–247 (2014).

11. C. L. Chen et al., Ascending neurons convey behavioural state to integrative sensory and action selection brain regions. Nat Neurosci 26, 682–695 (2023).

12. T. Akam et al., The Anterior Cingulate Cortex Predicts Future States to Mediate Model-Based Action Selection. Neuron 109, 149–163 e147 (2021).

13. A. W. Corcoran, J. Hohwy, in The Interoceptive Mind: From Homeostasis to Awareness, M. Tsakiris, H. De Preester, Eds. (Oxford University Press, 2018), pp. 0.

14. C. Gizowski, C. Zaelzer, C. W. Bourque, Clock-driven vasopressin neurotransmission mediates anticipatory thirst prior to sleep. Nature 537, 685–688 (2016).

15. V. Augustine, S. K. Gokce, Y. Oka, Peripheral and Central Nutrient Sensing Underlying Appetite Regulation. Trends Neurosci 41, 526–539 (2018).

16. Y. Livneh et al., Estimation of Current and Future Physiological States in Insular Cortex. Neuron 105, 1094–1111 e1010 (2020).

17. C. Deans, Biological Prescience: The Role of Anticipation in Organismal Processes. Front Physiol 12, 672457 (2021).

18. B. S. McEwen, J. C. Wingfield, The concept of allostasis in biology and biomedicine. Horm Behav 43, 2–15 (2003).

19. W. E. Allen et al., Thirst regulates motivated behaviour through modulation of brainwide neural population dynamics. Science 364, 253-+ (2019).

20. R. Gong, S. J. Xu, A. Hermundstad, Y. Yu, S. M. Sternson, Hindbrain Double-Negative Feedback Mediates Palatability-Guided Food and Water Consumption. Cell 182, 1589-+ (2020).

21. Y. Mandelblat-Cerf et al., Arcuate hypothalamic AgRP and putative POMC neurons show opposite changes in spiking across multiple timescales. Elife 4, (2015).

22. R. M. Ryan, Psychological Needs and the Facilitation of Integrative Processes. J Pers 63, 397–427 (1995).

23. Y. M. Chen, Y. C. Lin, T. W. Kuo, Z. A. Knight, Sensory Detection of Food Rapidly Modulates Arcuate Feeding Circuits. Cell 160, 829–841 (2015).

24. C. J. Burnett et al., Hunger-Driven Motivational State Competition. Neuron 92, 187–201 (2016).

25. S. M. Sternson, A. K. Eiselt, Three Pillars for the Neural Control of Appetite. Annu Rev Physiol 79, 401–423 (2017).

26. G. G. Berntson, S. S. Khalsa, Neural Circuits of Interoception. Trends in Neurosciences 44, 17–28 (2021).

27. R. A. Hinde, Ethological Models and the Concept of Drive. Brit J Philos Sci 6, 321–331 (1956).

28. D. S. Lehrman, A Critique of Konrad Lorenz’s Theory of Instinctive Behaviour. The Quarterly Review of Biology 28, 337–363 (1953).

29. R. Dawkins, A threshold model of choice behaviour. (1969).

30. K. Z. Lorenz, The Comparative Method in Studying Innate Behaviour Patterns. Sym Soc Exp Biol 4, 221–268 (1950).

31. S. Luquet, F. A. Perez, T. S. Hnasko, R. D. Palmiter, NPY/AgRP neurons are essential for feeding in adult mice but can be ablated in neonates. Science 310, 683–685 (2005).

32. M. A. Cowley et al., Leptin activates anorexigenic POMC neurons through a neural network in the arcuate nucleus. Nature 411, 480–484 (2001).

33. Q. Gao, T. L. Horvath, Neurobiology of feeding and energy expenditure. Annu Rev Neurosci 30, 367–398 (2007).

34. T. M. Hahn, J. F. Breininger, D. G. Baskin, M. W. Schwartz, Coexpression of Agrp and NPY in fasting-activated hypothalamic neurons. Nature Neuroscience 1, 271–272 (1998).

35. J. D. Deem, C. L. Faber, G. J. Morton, AgRP neurons: Regulators of feeding, energy expenditure, and behaviour. Febs J 289, 2362–2381 (2022).

36. I. C. Alcantara, A. P. M. Tapia, Y. Aponte, M. J. Krashes, Acts of appetite: neural circuits governing the appetitive, consummatory, and terminating phases of feeding. Nat Metab 4, 836–847 (2022).

37. Q. Q. Liu et al., An iterative neural processing sequence orchestrates feeding. Neuron 111, (2023).

38. Y. H. Lee et al., Lateral hypothalamic leptin receptor neurons drive hunger-gated food-seeking and consummatory behaviours in male mice. Nat Commun 14, (2023).

39. S. Shin et al., Early adversity promotes binge-like eating habits by remodeling a leptin-responsive lateral hypothalamus-brainstem pathway. Nature Neuroscience 26, 79-+ (2023).

40. J. Bosulu, G. Pezzulo, S. Hétu, A computational account of needing and wanting. bioRxiv, 2022.2010.2024.513547 (2023).

41. O. J. Hulme, T. Morville, B. Gutkin, Neurocomputational theories of homeostatic control. Phys Life Rev 31, 214–232 (2019).

42. H. G. R. Kim et al., A Unified Framework for Dopamine Signals across Timescales. Cell 183, 1600-+ (2020).

43. J. Cox et al., A neural substrate of sex-dependent modulation of motivation. Nat Neurosci 26, 274–284 (2023).

44. K. Choi et al., Distributed processing for value-based choice by prelimbic circuits targeting anterior-posterior dorsal striatal subregions in male mice. Nat Commun 14, 1920 (2023).

45. S. Schneider, J. H. Lee, M. W. Mathis, Learnable latent embeddings for joint behavioural and neural analysis. Nature 617, 360–368 (2023).

46. S. Rutherford et al., The normative modeling framework for computational psychiatry. Nat Protoc 17, 1711–1734 (2022).

47. M. R. Zimmer, A. H. O. Fonseca, O. Iyilikci, R. D. Pra, M. O. Dietrich, Functional Ontogeny of Hypothalamic Agrp Neurons in Neonatal Mouse Behaviours. Cell 178, 44-+ (2019).

48. Y. M. Chen et al., Sustained NPY signaling enables AgRP neurons to drive feeding. Elife 8, (2019).

49. Y. Aponte, D. Atasoy, S. M. Sternson, AGRP neurons are sufficient to orchestrate feeding behaviour rapidly and without training. Nature Neuroscience 14, 351–355 (2011).

50. J. N. Siemian, M. A. Arenivar, S. Sarsfield, C. B. Borja, C. N. Russell, Y. Aponte, Lateral hypothalamic LEPR neurons drive appetitive but not consummatory behaviours. Cell Rep 36, 109615 (2021).

51. T. W. Chen et al., Ultrasensitive fluorescent proteins for imaging neuronal activity. Nature 499, 295–300 (2013).

52. H. Akaike, Citation Classic - a New Look at the Statistical-Model Identification. Cc/Eng Tech Appl Sci, 22–22 (1981).

53. J. J. O. de Xivry, P. Lefevre, A switching cost for motor planning. J Neurophysiol 116, 2857–2868 (2016).

54. F. M. Toates, Homeostasis and drinking. Behavioural and Brain Sciences 2, 95–136 (1979).

55. A. K. Seth, Interoceptive inference, emotion, and the embodied self. Trends Cogn Sci 17, 565–573 (2013).

56. M. A. Apps, M. Tsakiris, The free-energy self: a predictive coding account of self-recognition. Neurosci Biobehav Rev 41, 85–97 (2014).

57. M. Hovd, R. R. Bitmead, Feedforward for stabilization in the presence of constraints. J Process Contr 22, 659–665 (2012).

58. B. Senapati et al., A neural mechanism for deprivation state-specific expression of relevant memories in Drosophila. Nature Neuroscience 22, 2029-+ (2019).

